# A general thermodynamic approach for model reduction of enzyme cycles and electrogenic transporters

**DOI:** 10.64898/2026.06.16.732208

**Authors:** Michael Pan, Peter J. Gawthrop, Joseph Cursons, Edmund J. Crampin

## Abstract

Mathematical models of enzyme cycles form the basis of quantifying key features of metabolism and membrane transport. These models are often integrated into more comprehensive models such as whole-cell models to understand emergent behaviours between interacting components. However, it is currently computationally infeasible to simulate the full dynamical behaviour of every enzyme at a network scale. Model reduction is frequently used to improve computational efficiency, but in general, these approaches do not preserve physical and thermodynamic consistency.

Here, we outline a general method for simplifying enzyme kinetics models while retaining mass, charge and energy balance. We base our approach on the bond graph, which is a general methodology for modelling biological systems from fundamental physical laws. This approach ensures that key physical constraints are enforced in every model, regardless of their complexity. Our thermodynamic model reduction framework is readily extended to electrogenic transporters through the coupling of chemical and electrical processes. Through the application of our approach to both hypothetical enzyme cycles and real data from the Na^+^/K^+^ ATPase, we show that it can rapidly screen for plausible network structures in circumstances where enzyme catalytic mechanisms may not be fully characterised, facilitating biological discovery and drug development.

## 1 Introduction

Enzyme cycles are essential to cellular function, catalysing reactions that would otherwise proceed too slowly spontaneously. The activation or inactivation of an enzyme is a key mechanism that cells use to control the rate of a reaction, and thus direct metabolites along desired pathways [1]. The concept of enzyme cycles can also be generalised to multi-physical thermodynamic cycles, which govern many ion transport processes that are essential to signalling and maintaining cell volume [2]. Mathematical models of enzyme cycles have proven invaluable in interpreting experimental data, with the Michaelis-Menten model being widely used to characterise the turnover and affinity of enzymes using experimental assays [3]. As kinetic models of enzymes have become more detailed over time, they have been used in conjunction with experimental data to provide insights into potential mechanisms of enzyme catalysis [4].

Recently, there has been a rapidly growing interest in using enzyme cycle models as part of more comprehensive systems biology models, including whole-cell models that aim to dynamically simulate all known biomolecules over time. There are currently two published whole-cell models of *Mycoplasma genitalium* and the JCVI-syn3A minimal cell, with a further whole-cell model of *Escherichia coli* in active development [5–8]. However, there are some widely acknowledged bottlenecks in whole-cell modelling that apply to enzyme cycles [9]. One challenge is the substantial computational cost of simulating whole-cell models. On the scale of entire metabolic networks, it is infeasible to model the dynamics of each enzyme in full mechanistic detail. Thus, whole-cell models employ various approaches for reducing model complexity. The *M. genitalium* and *E. coli* whole-cell models represent metabolism using constraint-based models to simplify simulation [5, 7, 8], whereas Thornburg et al. [6] use a convenience kinetics equation to reduce the order of their metabolic models.

While the importance of computational efficiency is acknowledged in large-scale systems biology models, approaches to model reduction in general do not satisfy the requirement for physical and thermodynamic consistency [10, 11]. Many existing reaction networks break the laws of thermodynamics, through the use of either irreversible reactions or ill-defined parameters that violate detailed balance [10, 12–14]. Due to their reliance on constraint-based models with irreversible reactions, the whole-cell models of *M. genitalium* and *E. coli* are not thermodynamically consistent [5, 7, 8]. The JCVI-syn3A whole-cell model [6] resolves this issue by using convenience kinetics as a general equation structure for all enzymes [15, 16]. However, while it is relatively simple to implement and satisfies thermodynamic consistency, the convenience kinetics rate law assumes a rapid and random-order substrate binding mechanism, which is unsuitable for modelling certain enzymes [4]. In reality, enzymes employ a highly diverse range of reaction mechanisms, and it is unlikely that the catalytic mechanisms of all enzymes would be reducible to a small subset of rate laws. Thus, we anticipate that enzymatic rate laws would gradually need to be replaced as experimental data become more available for models to utilise.

Model calibration is an essential aspect of developing systems biology models. In the context of enzymes and transporters, model calibration has to date mostly focussed on parameter estimation and uncertainty quantification under the assumption of a known model structure [17, 18]. However, for many enzymes, we have limited data and incomplete knowledge about their mechanism. As a result, structural uncertainty and model selection have been relatively underexplored [19, 20]. One potential contributor to the relative lack of work in this area is the time-consuming process of manually deriving rate laws by hand. Therefore, we believe that automating this process would make techniques such as model selection more accessible.

In this study, we address the above limitations by formulating a model reduction approach that is generalisable across enzyme cycles while preserving thermodynamic consistency. As this approach is computable, it reduces the time required to develop such reduced enzyme kinetic models. The bond graph approach was originally developed for modelling engineering systems using energy flows [21–24]. In particular, the approach can be used for energy-based model reduction [21]. For this reason, we base our approach on the bond graph, which has been shown to provide a general and powerful methodology for modelling biological systems [12, 25–30]. This approach encodes both the fundamental biophysical laws governing model components and conservation laws associated with network topology, and has been used to model biological systems from enzymes to ion transport and mechanobiology [12, 31–33]. An additional advantage of bond graphs is that they are inherently modular, allowing submodules to be developed in isolation before coupling these into composite models [34–36].

At the core of our strategy is a general method for simplifying models of enzymes, combining the use of bond graphs with timescale separation approaches such as rapid equilibrium and the quasi-steady-state approximation [37]. The use of simplified rate laws rather than differential equations makes parameter estimation computationally efficient. Because our method is general for a wide range of enzyme cycles, it is easily implemented in software. Thus, our approach can be used to test the validity of several candidate mechanisms against an experimental dataset to assess validity. Our methodology enables the development of computationally efficient, biochemically plausible and thermodynamically consistent models that are also reusable and modular.

In Subsection 2.1.1, we review the essential concepts of enzyme model reduction using a simple reversible Michaelis-Menten model. We then motivate the use of an energetic framework for representing enzyme kinetics (Subsection 2.1.2–2.1.3), and outline a generalised methodology for reducing models of enzyme kinetics under this energetic framework (Subsection 2.2). In Subsection 3.1, we verify that our approach can be used to generate thermodynamically consistent rate laws for a range of enzyme cycles. Finally, we show how results can be compared between multiple potential mechanisms to systematically compare candidate models to data, and apply this approach to two examples: a hypothetical enzyme with a bi-bi mechanism in which substrates bind individually in an ordered sequence before the products are produced and sequentially released (Subsection 3.2); and a physiological example in the cardiac Na^+^/K^+^ ATPase (Subsection 3.3).

## 2 Methods

### 2.1 Reversible Michaelis-Menten enzyme

#### 2.1.1 Kinetic formulation

We first review the essential principles of model reduction that we use in our approach. A simple Michaelis-Menten model (Figure 1) is used to illustrate how these principles can be applied to enzyme kinetics. Because all enzymes are thermodynamically reversible, we differ from the typical formulation in using a reversible second reaction (Figure 1A). Using the law of mass action, the rates of the two reactions are

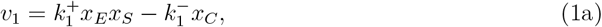

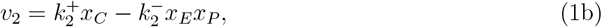

and accumulation of free enzyme (E) and complex (C) are given by the ordinary differential equations (ODEs)

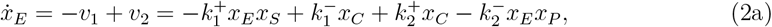

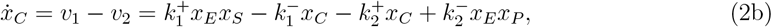

whre 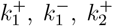 and 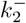 are the kinetic constants, and *x*_*s*_ is the molar amount of species *s∈* {E, C, S, P} . Here we assume that all of the species exist within the same volume so that their molar amounts are proportional to their concentration.

**Figure 1:**
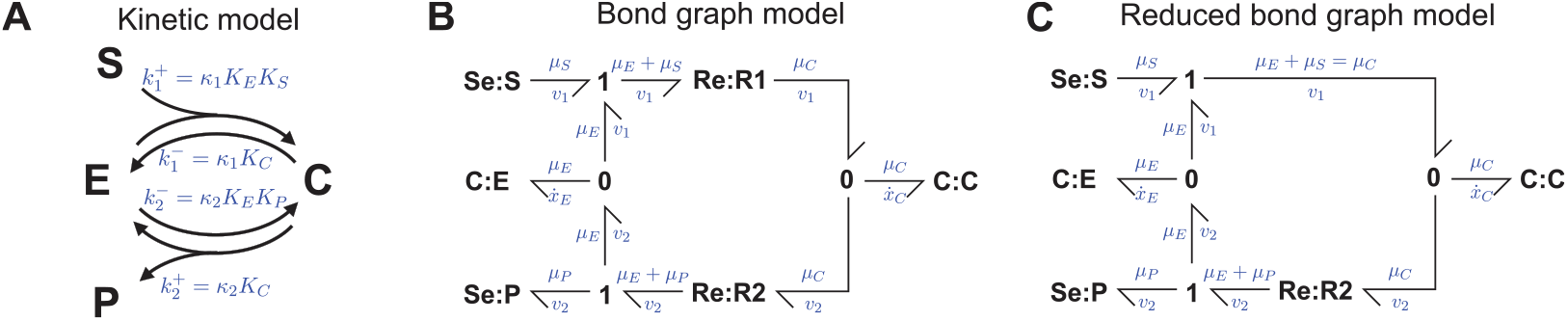
A reversible Michaelis-Menten model. **(A)** Kinetic representation, with the mapping between kinetic and bond graph parameters shown in blue; **(B)** Bond graph representation of the full model; **(C)** Bond graph representation of the model simplified using rapid equilibrium; the **Re:R1** component is removed to enforce equilibrium between the potentials *µ*_*E*_ + *µ*_*S*_ and *µ*_*C*_.

In the context of cells, enzymes are typically lower in concentration than their substrates [10]. We therefore assume that the dynamics of the metabolites are slow relative to the dynamics of the enzyme, so that the enzyme is operating under a quasi-steady-state. Under this assumption – the quasi-steady-state approximation – the enzyme is at a steady-state for any given concentration of substrate (S) and product (P) [38]. Thus, the left hand side of Eq. 2 is effectively zero:

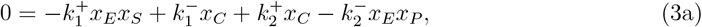

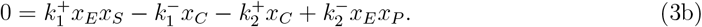

By solving the equations simultaneously, the amounts of enzyme and complex are

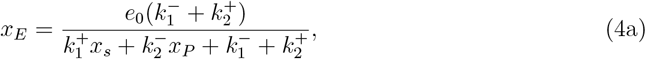

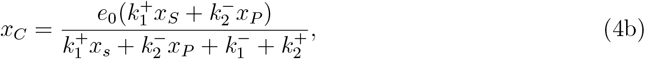

where

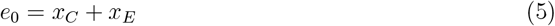

is the total amount of enzyme. Therefore, by substituting into Eq. 1a, the enzymatic reaction rate is

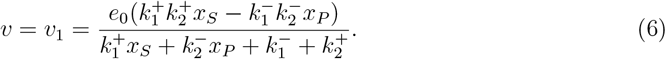

A key advantage of the quasi-steady-state approximation is that the individual enzyme states E and C are lumped into a total enzyme amount *e*_0_, and thus reaction rates can be calculated without simulating the dynamics of each conformation [39]. Because the steady-state cycling velocity is proportional to *e*_0_ [39, 40], it is useful to define its normalised form *v*_cyc_ = *v/e*_0_ as the cycling rate [s^*−*1^] for use in later sections.

Rate laws can also be simplified by assuming that some steps occur significantly faster than others so that they are effectively at equilibrium. This is commonly applied to the Michaelis-Menten model through the rapid equilibrium approximation, where the first reaction operates on a faster timescale relative to the second reaction and is therefore at equilibrium. In this approximation, the forward and reverse fluxes of the first reaction are equal:

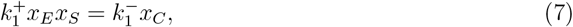

so the total amount of enzyme can be expressed as

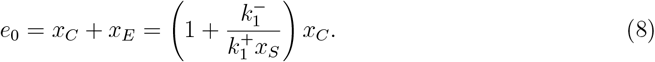

By substituting this into Eq. 1b, it can be shown that the reaction rate is

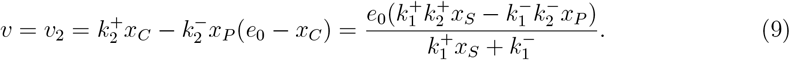

We note that the rapid equilibrium approximation is distinct from the quasi-steady-state approximation; neither approximation implies the other, and they can be applied separately or in conjunction with each other. The quasi-steady-state approximation is able to reduce enzyme kinetics to a single rate law, but does not reduce the number of parameters. By contrast, the rapid equilibrium approximation reduces the number of parameters but does not reduce the kinetics of enzymes to a single rate law. Thus, these two approaches are often used in conjunction with one another to harness the strengths of both [37].

#### 2.1.2 Energetic formulation

While the kinetic formulation can be used to apply model reduction to enzyme kinetics, a limitation of this formulation is that kinetic parameters are not independent, as they contain both species and reaction-related properties. As a result, these parameters cannot be perturbed individually, and modellers frequently need to apply additional detailed balance constraints to satisfy thermodynamic consistency [14]. Here we describe an alternative formulation, the energetic formulation, which separates thermodynamic quantities from kinetic quantities [12, 14– 16, 41]. Accordingly, species and reactions are separate components with independent parameters. Each species *s* is associated with a chemical potential *µ*_*s*_ [J*/*mol], described by the equation

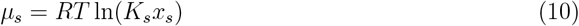

where *R* = 8.314 J*/*mol*/*K is the universal gas constant, *T* [K] is the absolute temperature, *K*_*s*_ [mol^*−*1^] is the species thermodynamic constant, and *x*_*s*_ [mol] is the molar amount of species. This chemical potential describes the propensity of the chemical species to drive reactions. Reactions are dissipative processes that operate in the direction of decreasing chemical potential. In the energetic formalism, a reaction *r* is described by the Marcelin-de Donder equation

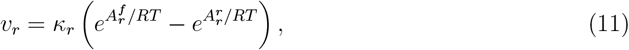

where *v*_*r*_ [mol/s] is the rate of reaction (which can be positive or negative), *κ*_*r*_ [mol/s] is the reaction rate constant, 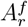 [J/mol] is the forward affinity (the sum of chemical potentials of the reactants), and 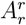 [J/mol] is the reverse affinity (the sum of chemical potentials of the products).

In the Michaelis-Menten model, the chemical potentials are

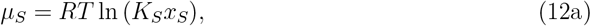

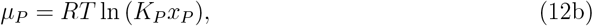

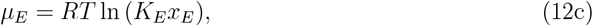

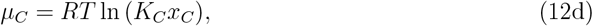

and the reaction affinities are

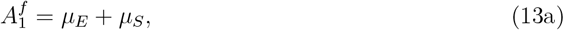

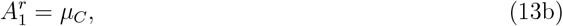

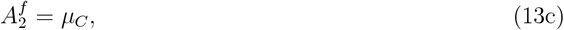

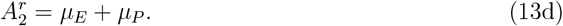

Using Eqs. 11–13, it can be shown that the reaction rates of the full system are

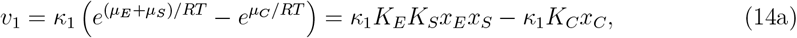

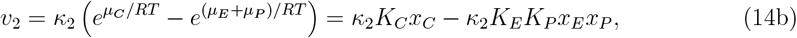

and the differential equations for the rates of change for the enzyme states are as described in Eq. 2, with the kinetic relations

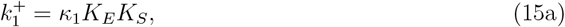

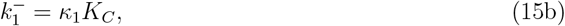

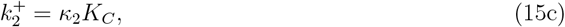

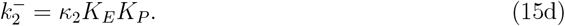

It should be noted that while the energetic parameters *K*_*s*_ and *κ*_*r*_ give rise to the same kinetic behaviour as the kinetic constants, they also determine the chemical potentials associated with each species and reaction. Because energetic parameters are thermodynamically independent, they are guaranteed to be thermodynamically consistent and can be perturbed or updated independently [14–16, 42].

The analysis for the quasi-steady-state approximation follows that for the kinetic formulation with the substitutions in Eq. 15, therefore the rate of the catalysed reaction is

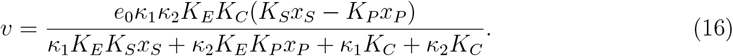

With some algebraic manipulation, Eq. 16 is equivalent to Eq. 5.16 of Gawthrop and Crampin [12], which is an equation for the reaction rate derived from the same system.

The analysis for the rapid equilibrium approximation also follows a similar pattern to the kinetic framework. However, it is useful to note that the rapid equilibrium constraint in Eq. 7 arises due to an equilibration of affinities on both sides of reaction 1:

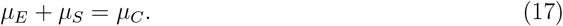

Therefore, by using Eq. 12, the rapid equilibrium constraint can be alternatively formulated as

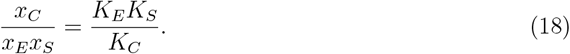

This constraint can be used to show that the reaction rate is

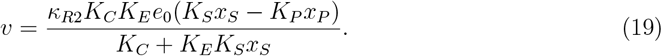

We note that Eq. 19 is Eq. 16 when we take *κ*_1_ → *∞*. Thus, an advantage of formulating rapid equilibrium in terms of energetic parameters is that the approximation can be linked back to the parameter for the corresponding reaction.

#### 2.1.3 Bond graph representation

The thermodynamic formulation in Subsection 2.1.2 is generalised by encoding it as a bond graph. A bond graph representation of the reversible Michaelis-Menten model is shown in Figure 1B. In this representation, components are linked by bonds (*⇁*) that represent the transfer of power. For biochemical systems, the power carried by each bond is decomposed into a chemical potential *µ* [J/mol] and a molar flow rate *v* [mol/s] which multiply to give power *P* = *µv*. Because bond graphs explicitly model energy transfer, thermodynamic consistency is automatically satisfied along with many other conservation laws [21, 23, 43]. The quantities carried by each bond are constrained by the components they are connected to. These constraints dictate the behaviour of the system. The constraints for the Michaelis-Menten model (Eqs. 2,10–14) are captured using **C, Se, Re, 0** and **1** components.

Dynamic chemical species are represented using **C** components, which are energy storage components that are named as such due to their analogy with electrical capacitors. There are two **C** components in the bond graph representation corresponding to the dynamic species E and C. **C** components encode the constraints in Eqs. 12c–12d. The two **C** components are dynamic as the molar amounts of species are allowed to vary based on the molar flow rates of their connected bonds.

The metabolites S and P are represented using **Se** components. **Se** components relate chemical potential and molar amount in the same way as **C** components (Eqs. 12a–12b). However, in contrast to the enzyme states, concentrations of metabolites are assumed to be constant on the timescale of enzyme kinetics. **Se** components represent this behaviour by fixing the concentrations of these metabolites to constant values 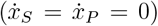. These species with fixed concentrations are known as *chemostats*. Because external flows are required to keep the concentrations of chemostats constant, chemostats represent connections to external energy sources or sinks. When incorporating enzyme kinetics models into broader whole-cell models with variable metabolite concentrations, external connections can be formalised in terms of **SS** components [42].

Reactions are represented using **Re** components. There are two **Re** components in Figure 1B corresponding to the two reactions of the Michaelis-Menten model, and these encode Eq. 11. Because reactions dissipate energy, they are analogous to electrical resistors.

Conservation laws are represented using **0** and **1** junctions. The **0** junction is used when a species is involved in multiple reactions. This junction encodes the conservation of mass law seen in the first equality of Eq. 2, and also ensures that the chemical potential contributed by the species to each reaction is equal (see chemical potentials connected to the **0** junction in Figure 1B). Therefore the **0** junction is a generalisation of Kirchhoff’s current law. Similarly, **1** junctions are used when more than one species is present on either side of a reaction. This junction encodes the conservation of potential law in Eqs. 13a,13d while ensuring that the molar flow rates of the connected bonds are equal (see Figure 1B). Thus, the **1** junction is a generalisation of Kirchhoff’s voltage law. Via the analysis in Subsection 2.1.2, the constraints encoded by the bond graph components are sufficient to derive the dynamics of the Michaelis-Menten model.

Because the bond graph representation is energetic, it can naturally represent the rapid equilibrium approximation. As seen in Eq. 17, rapid equilibrium is the equilibration of the forward and reverse affinities. Bond graphs can represent this constraint through the removal of the corresponding **Re** component (Figure 1C) [12].

For a more comprehensive description of bond graph theory, the reader is referred to the textbooks by Gawthrop and Smith [21] and Borutzky [43]. The application of bond graphs to biochemical systems is detailed in Gawthrop and Crampin [12], Gawthrop, Cursons and Crampin [42], Pan et al. [44] and Pan et al. [34].

### 2.2 General model reduction methodology

To extend our analysis to arbitrarily large enzyme cycles, we use timescale separation and stoichiometric matrices [45] to derive explicit equations for the cycling rate. Because these equations are derived explicitly, they are easily generated using software and scalable to complex enzyme models. We briefly summarise the method here, with the full method described in Appendix A of the electronic supplementary material.

As seen in Subsection 2.1.1, the derivation of a rate law involves three constraints corresponding to the rapid equilibrium approximation (Eq. 18), quasi-steady-state approximation (Eq. 3), and the conservation law arising from conservation of enzyme (Eq. 5). The *rapid equilibrium* constraint arises from the steady state (and also thermodynamic equilibrium) of the fast timescale:

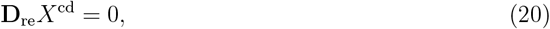

while the *quasi-steady-state* constraint arises from the steady state of the slow timescale:

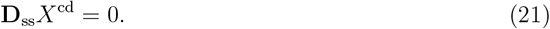

*X*^cd^ is a vector containing the amounts of each enzyme state, and **D**_re_ and **D**_ss_ are matrices that depend on the energetic parameters, metabolite concentrations and the stoichiometry of reactions within the enzyme cycle. Eqs. 20 and 21 are derived in Appendix A of the electronic supplementary material, and their full forms are given in Eqs. S10 and S18 respectively. Note that for computational efficiency, the rapid equilibrium constraint is derived prior to deriving the quasi-steady-state constraint to reduce the dimensionality of the equations. The steady state of the system is dependent on the amount of enzyme present in the initial condition of the system:

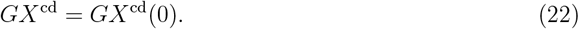

For the enzyme cycles examined in this study, *G* is a row vector of ones, and *e*_0_ = *GX*^cd^(0) is the total amount of enzyme. Because **D**_re_, **D**_ss_ and *G* are independent of *X*^cd^, linear methods can be used to simultaneously solve Eqs. 20–22 for *X*^cd^, which can then be substituted into the appropriate Marcelin-de Donder equation (Eq. 11) to derive an analytical expression for the cycling rate.

In the Michaelis-Menten example with the rapid equilibrium approximation (analysis detailed in Subsection 2.1.2 with bond graph in Figure 1C), the matrix equations are

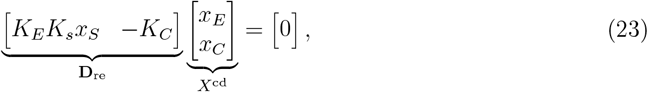

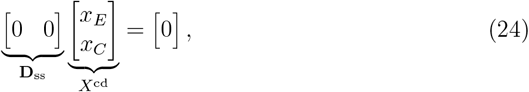

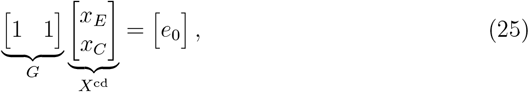

and thus the enzyme states can be solved through the matrix equation

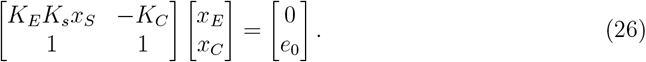

where Eq. 24 is removed as it provides no additional information.

Similarly, for the Michaelis-Menten model with the quasi-steady-state approximation, there is no rapid equilibrium constraint and

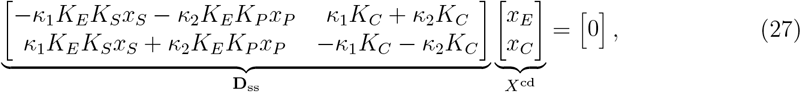

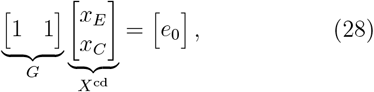

and the enzyme states are solved through the equation

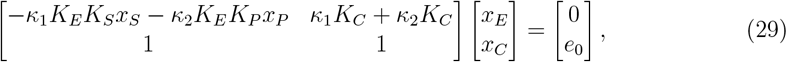

where the second row of **D**_ss_ is omitted due to its linear dependency with the first row.

### 2.3 Parameter estimation

Energetic parameters are estimated by minimising a cost function. The cost function is minimised using the TikTak multistart algorithm [46] with the Limited-memory Broyden-Fletcher-Goldfarb-Shanno algorithm as the local optimiser [47]. The TikTak algorithm is initialised with 1000 random starting points.

Energetic parameters describe the independent contribution of each species to equilibrium constants. As a result, there are families of energetic parameters that give rise to the same kinetic behaviour [42, 44] and therefore families of parameter values rather than specific parameter values arise from fitting kinetic measurements [16, 44]. By analysing the nullspaces of the stoichiometric matrix [42, 45], we develop a fitting procedure that expresses the resulting parameters as mathematical expressions with free parameters that can be assigned arbitrarily (Appendix B of the electronic supplementary material). The energetic parameters associated with species and reactions are positive, therefore they are log transformed during optimisation for computational efficiency [48].

A squared residuals cost function is used for parameterising the models in Subsections 3.1–3.2. For the Na^+^/K^+^ ATPase (Subsection 3.3), the weighted squared residuals cost function described in [44] is used:

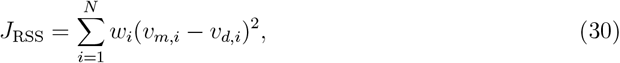

where *i* = 1, 2, …, *N* is the index for each experimental measurement, *v*_*m,i*_ is the model prediction of cycling rate and *v*_*d,i*_ is the measured cycling rate. We add an additional cost

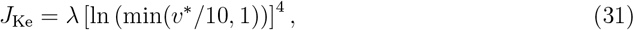

where *λ* = 10^6^ and *v*^*\**^ is the (non-normalised) cycling rate at [K^+^]_e_ = 5.4 mM. This term penalises low cycling rates and is used to ensure that the magnitudes of cycling rates are within realistic ranges for the Nakao and Gadsby [49] extracellular K^+^ dataset. Thus, the objective function used by the optimiser is

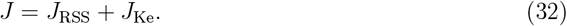

### 2.4 Software implementation

A Julia implementation of the bond graph based model reduction approach is available at https://github.com/mic-pan/EnzymeModelIdentification.jl. The `EnzymeModelIdentification.jl` package supports the automatic generation of cycling rates from reaction networks corresponding to enzyme catalysed reactions. Symbolic model reduction is performed using the `Catalyst.jl` and `SymbolicUtils.jl` packages. Users can load in experimental data via the `DataFrames.jl` package and fit model parameters via an interface to `Optimization.jl. EnzymeModelIdentification.jl` also supports the reduction and calibration of electrogenic transporter models.

## 3 Results

### 3.1 Model reduction in bi-bi enzyme catalysed reactions

#### 3.1.1 Reduced model equations

To demonstrate that our method is generalisable, we apply it to the more complex four-state enzyme cycle shown in Figure 2A, which represents an ordered bi-bi mechanism [37]. Reactions 1 and 4 (enclosed in dotted blue boxes) are assumed to be in rapid equilibrium. The bond graph of the reaction network is shown in Figure 2B and the system with rapid equilibrium applied is shown in Figure 2C. Using the method outlined in Subsection 2.2, the cycling rate of this enzyme is given by

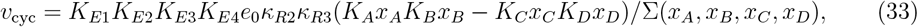

where

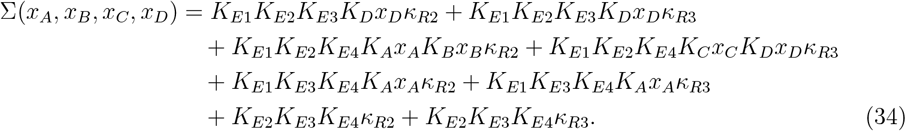

**Figure 2:**
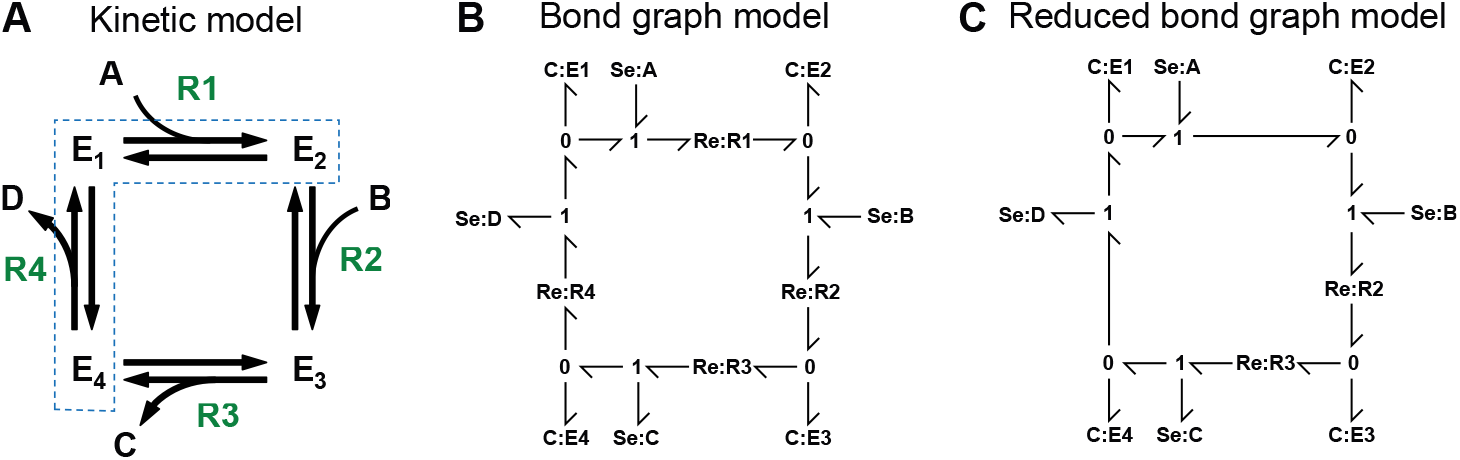
A four-state enzyme cycle. **(A)** Kinetic representation, with the reaction names shown in green. Reactions assumed to be in rapid equilibrium are enclosed in the dotted blue box. **(B)** Bond graph representation of the full model; **(C)** Bond graph representation of the model with the **Re:R1** and **Re:R4** components are removed to enforce equilibrium between the potentials *µ*_*E*1_ + *µ*_*A*_ and *µ*_*E*2_ and between the potentials *µ*_*E*4_ + *µ*_*C*_ and *µ*_*E*1_.

To verify that the cycling rate (Eqs. 33–34) of the four-state model exhibits the expected limiting behaviour as the rates of reactions 1 and 4 increase, we plot the cycling rate for the full model with only quasi-steady-state approximation applied, and the model approximated by rapid equilibrium (Figure 3A). We use the parameters *κ*_2_ = *κ*_3_ = 1, *K*_*E*1_ = 4, *K*_*E*2_ = 2, *K*_*E*3_ = 1, *K*_*E*4_ = 2, *K*_*A*_ = 2, *K*_*B*_ = 1, *K*_*C*_ = 1 and *K*_*D*_ = 2 and set the rates of reactions 1 and 4 to the same value κ that we vary. As κ increases, the magnitude of the flux increases in both the forward and reverse modes of operation, and the curve approaches that of the rapid equilibrium approximation, as expected.

**Figure 3:**
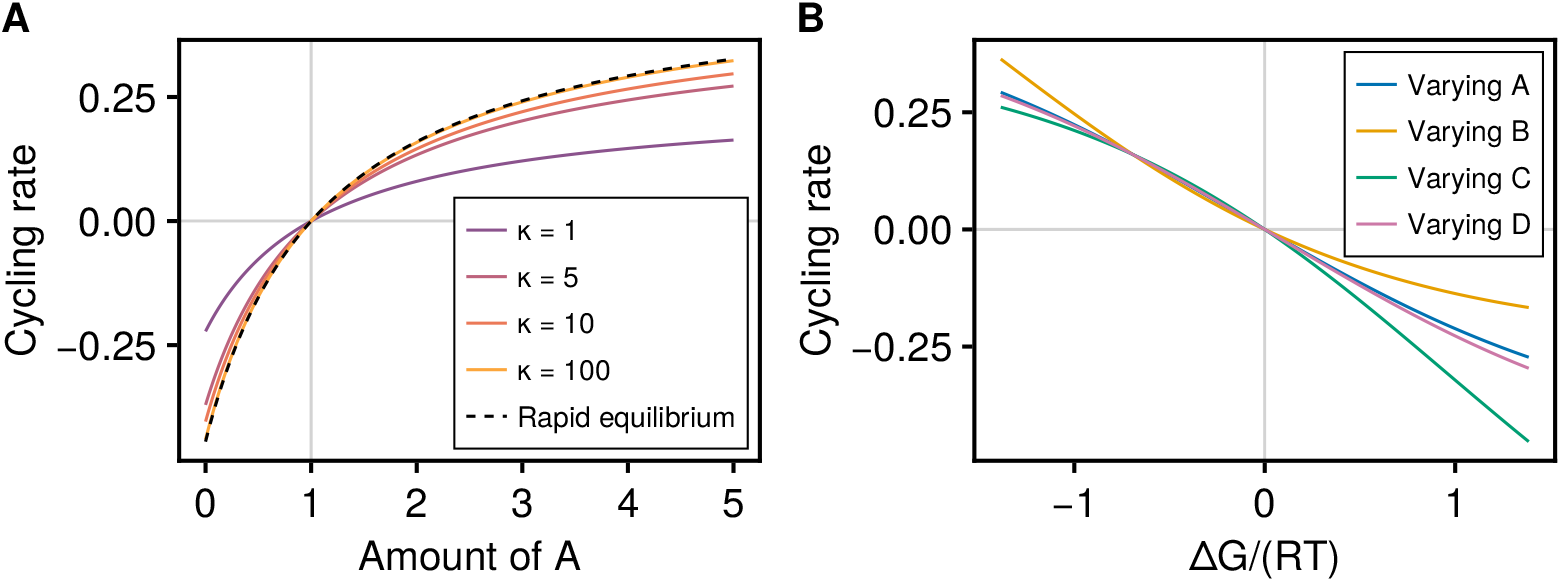
The cycling rate of the four-state enzyme model. **(A)** Comparison of cycling rates of the full model with different values of *κ*, when compared to the model with the rapid equilibrium approximation. **(B)** The relationship between Gibbs free energy and cycling rate when different metabolites are varied to change the free energy. Metabolite amounts are varied from the base conditions (*x*_*A*_, *x*_*B*_, *x*_*C*_, *x*_*D*_) = (2, 1, 1, 1).

#### 3.1.2 Thermodynamic consistency

Because the model is specified within an energetic formalism, the simplified rate law is thermodynamically consistent. Eq. 33 contains the term (*K*_*A*_*x*_*A*_*K*_*B*_*x*_*B*_ *− K*_*C*_*x*_*C*_*K*_*D*_*x*_*D*_) in the numerator, which ensures that the cycling rate is thermodynamically consistent. The free energy of the overall reaction *A* + *B* ⇌ *C* + *D* is

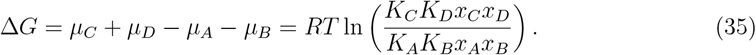

At equilibrium, Δ*G* = 0, hence the metabolites must satisfy the constraint

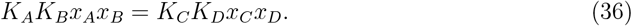

This is illustrated in Figure 3A, where regardless of whether the full or reduced model is used, the cycling rate is zero at the equilibrium point

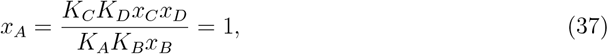

which corresponds to the thermodynamic equilibrium described by Eq. 36.

A more stringent check of the thermodynamic consistency of the system is to plot the Gibbs free energy of the overall reaction against cycling rate under several conditions (Figure 3B). Since Δ*G* can be changed by varying any of the metabolites, four curves are generated by varying each individually from a non-equilibrium regime of operation where (*x*_*A*_, *x*_*B*_, *x*_*C*_.*x*_*D*_) = (2, 1, 1, 1). While varying each of the metabolites results in different kinetic behaviour, all curves pass through the equilibrium point Δ*G* = 0, *v*_cyc_ = 0, showing that the system is thermodynamically consistent under a wide range of conditions.

### 3.2 Benchmarking of alternative enzyme catalytic mechanisms against synthetic data

We next use synthetic enzyme assay data to investigate whether it is possible to identify a true enzyme catalytic mechanism from a set of candidate models. We consider the family of cyclic bi-bi enzyme catalytic mechanisms with two to four states, as detailed in Table 1. We assume that the true model has parameter values κ_*R*1_ = 1.2, κ_*R*2_ = 1.4, κ_*R*3_ = 1.6, κ_*R*1_ = 1.8, *K*_*E*1_ = 1.0, *K*_*E*2_ = 1.2, *K*_*E*3_ = 1.4 and *K*_*E*4_ = 1.6. Additionally, the substrate parameters are *K*_*A*_ = *K*_*B*_ = *K*_*C*_ = *K*_*D*_ = 1, which are assumed to be known and therefore fixed during parameter estimation. To simulate an enzyme assay, we generate synthetic data for the concentrations (*x*_*A*_, *x*_*B*_, *x*_*C*_, *x*_*D*_) *∈ {*1, 2, 3, 4, 5*}*^4^, resulting in 625 steady-state measurements.

**Table 1:** Enzyme-catalysed reaction schemes bi-bi enzyme catalytic mechanisms with the overall reaction *A* + *B* ⇌ *C* + *D*.

| Index | Reaction 1 | Reaction 2 | Reaction 3 | Reaction 4 |
| --- | --- | --- | --- | --- |
| 1 | $E_1 + A \rightleftharpoons E_2$ | $E_2 + B \rightleftharpoons E_3$ | $E_3 \rightleftharpoons E_4 + C$ | $E_4 \rightleftharpoons E_1 + D$ |
| 2 | $E_1 + A \rightleftharpoons E_2$ | $E_2 + B \rightleftharpoons E_3$ | $E_3 \rightleftharpoons E_4 + D$ | $E_4 \rightleftharpoons E_1 + C$ |
| 3 | $E_1 + A \rightleftharpoons E_2$ | $E_2 \rightleftharpoons E_3 + C$ | $E_3 + B \rightleftharpoons E_4$ | $E_4 \rightleftharpoons E_1 + D$ |
| 4 | $E_1 + A \rightleftharpoons E_2$ | $E_2 \rightleftharpoons E_3 + C$ | $E_3 \rightleftharpoons E_4 + D$ | $E_4 + B \rightleftharpoons E_1$ |
| 5 | $E_1 + A \rightleftharpoons E_2$ | $E_2 \rightleftharpoons E_3 + D$ | $E_3 + B \rightleftharpoons E_4$ | $E_4 \rightleftharpoons E_1 + C$ |
| 6 | $E_1 + A \rightleftharpoons E_2$ | $E_2 \rightleftharpoons E_3 + D$ | $E_3 \rightleftharpoons E_4 + C$ | $E_4 + B \rightleftharpoons E_1$ |
| 7 | $E_1 + A + B \rightleftharpoons E_2$ | $E_2 \rightleftharpoons E_3 + C$ | $E_3 \rightleftharpoons E_1 + D$ | — |
| 8 | $E_1 + A + B \rightleftharpoons E_2$ | $E_2 \rightleftharpoons E_3 + D$ | $E_3 \rightleftharpoons E_1 + C$ | — |
| 9 | $E_1 + A \rightleftharpoons E_2 + C$ | $E_2 + B \rightleftharpoons E_3$ | $E_3 \rightleftharpoons E_1 + D$ | — |
| 10 | $E_1 + A \rightleftharpoons E_2 + C$ | $E_2 \rightleftharpoons E_3 + D$ | $E_3 + B \rightleftharpoons E_1$ | — |
| 11 | $E_1 + A \rightleftharpoons E_2 + D$ | $E_2 + B \rightleftharpoons E_3$ | $E_3 \rightleftharpoons E_1 + C$ | — |
| 12 | $E_1 + A \rightleftharpoons E_2 + D$ | $E_2 \rightleftharpoons E_3 + C$ | $E_3 + B \rightleftharpoons E_1$ | — |
| 13 | $E_1 + A \rightleftharpoons E_2$ | $E_2 + B \rightleftharpoons E_3 + C$ | $E_3 \rightleftharpoons E_1 + D$ | — |
| 14 | $E_1 + A \rightleftharpoons E_2$ | $E_2 + B \rightleftharpoons E_3 + D$ | $E_3 \rightleftharpoons E_1 + C$ | — |
| 15 | $E_1 + A \rightleftharpoons E_2$ | $E_2 \rightleftharpoons E_3 + C + D$ | $E_3 + B \rightleftharpoons E_1$ | — |
| 16 | $E_1 + A \rightleftharpoons E_2$ | $E_2 + B \rightleftharpoons E_3$ | $E_3 \rightleftharpoons E_1 + C + D$ | — |
| 17 | $E_1 + A \rightleftharpoons E_2$ | $E_2 \rightleftharpoons E_3 + C$ | $E_3 + B \rightleftharpoons E_1 + D$ | — |
| 18 | $E_1 + A \rightleftharpoons E_2$ | $E_2 \rightleftharpoons E_3 + D$ | $E_3 + B \rightleftharpoons E_1 + C$ | — |
| 19 | $E_1 + A + B \rightleftharpoons E_2$ | $E_2 \rightleftharpoons E_1 + C + D$ | — | — |
| 20 | $E_1 + A \rightleftharpoons E_2 + C$ | $E_2 + B \rightleftharpoons E_1 + D$ | — | — |
| 21 | $E_1 + A \rightleftharpoons E_2 + D$ | $E_2 + B \rightleftharpoons E_1 + C$ | — | — |
| 22 | $E_1 + A + B \rightleftharpoons E_2 + C$ | $E_2 \rightleftharpoons E_1 + D$ | — | — |
| 23 | $E_1 + A + B \rightleftharpoons E_2 + D$ | $E_2 \rightleftharpoons E_1 + C$ | — | — |
| 24 | $E_1 + A \rightleftharpoons E_2 + C + D$ | $E_2 + B \rightleftharpoons E_1$ | — | — |
| 25 | $E_1 + A \rightleftharpoons E_2$ | $E_2 + B \rightleftharpoons E_1 + C + D$ | — | — |

Figure 4B compares the quality of fit between models when Model 1 (a sequential binding/unbinding model; Figure 4A) is used as the ground truth. Model 1 is the only model with a perfect fit to synthetic data generated from itself, suggesting that it may be possible to identify the correct reaction structure in principle. Nonetheless, when cycling rate is plotted against *x*_*A*_ for a subset of the models (Figure 4C), Model 2 provides an excellent fit to the synthetic data and Models 6 and 8 are qualitatively similar, with Model 20 providing a poor fit. Thus, under certain conditions, it may be possible to approximate the four-reaction model with a three-reaction model such as Model 8. Under the presence of Gaussian measurement noise with standard deviation 0.01, there is a general increase in prediction error, defined as the residual sum of squares (RSS) across all models (Figure S4A of the electronic supplementary material).

**Figure 4:**
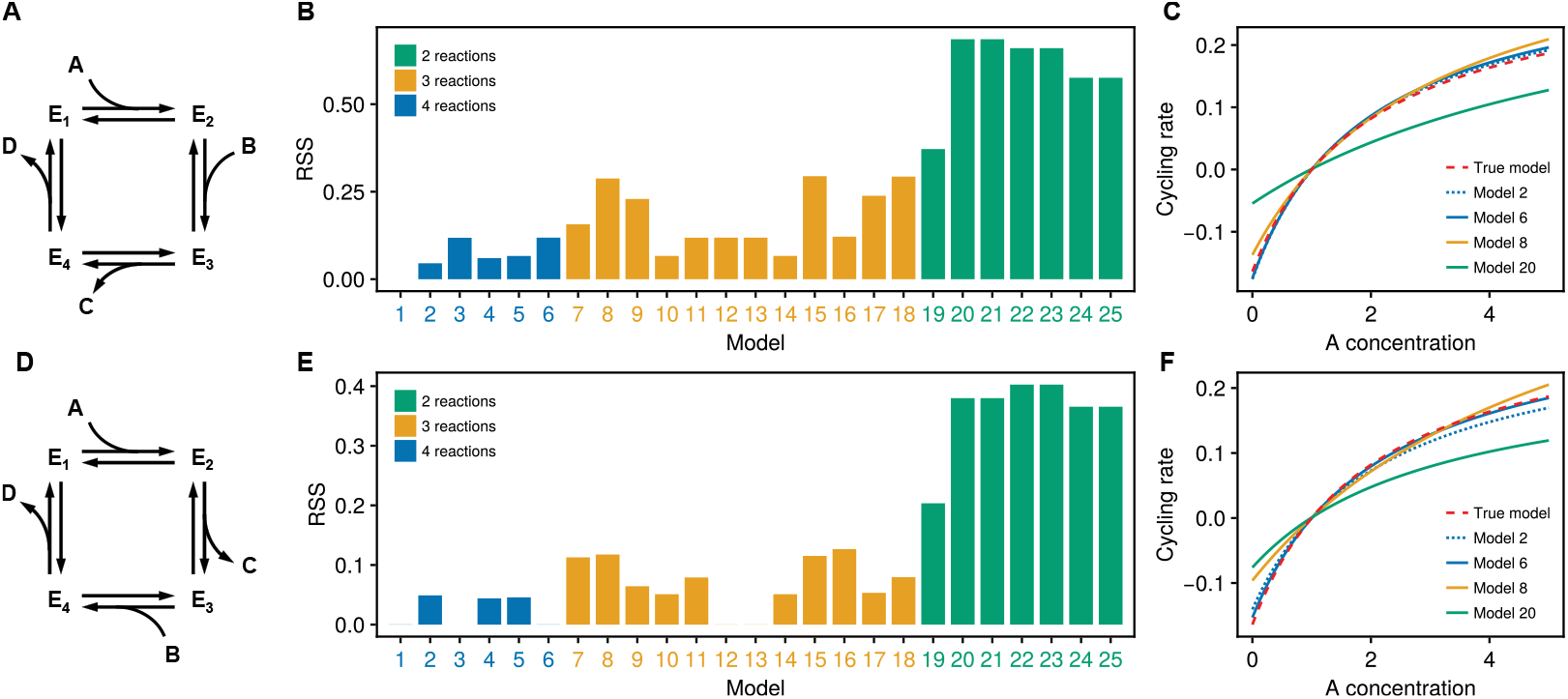
A comparison of fits of bi-bi enzyme catalytic mechanisms generated from synthetic data. Each row shows a comparison of model fits when synthetic data is generated from the model depicted on the left, with the top row corresponding to (A) Model 1 and the bottom row corresponding to (D) Model 3. Models are defined in Table 1. The middle columns (B) and (E) show the residual sum of squares (RSS) resulting from fitting each model to the synthetic data. (C) and (F) show a comparison of cycling rate curves for a representative subset of models. The cycling rate is plotted against *x*_*A*_, which is varied from 0 to 5.

When Model 3 (a ping-pong catalytic mechanism; Figure 4D) is used as the ground truth, Models 1, 6, 12 and 13 provide near perfect fits, even though only Model 3 results in a perfect fit with a residual sum of squares of zero (Figure 4E). This suggests that it would be very difficult to correctly identify Model 3 as the correct mechanism with the generated synthetic data, with the cycling rate plot in Figure 4F confirming that Model 6 is almost visually indistinguishable from Model 3. Thus, the presence of any measurement noise would make these models indistingushable given the data provided (see Figure S4B of the electronic supplementary material for fits with added noise). Interestingly, two sequential mechanisms (Models 1 and 6) are among those with near perfect fits, indicating that it may not even be possible to determine whether the mechanism is sequential or ping-pong in certain scenarios.

### 3.3 Application of model reduction to the cardiac Na^+^/K^+^ ATPase

The results of the previous section suggest that there is a degree of mechanistic uncertainty associated with fitting models to steady-state measurements that are typically gathered in enzyme assays [50]. A practical choice for dealing with this uncertainty is to select the simplest model that fits the data well, as simple models are less prone to over-fitting [51]. We applied this approach to a physiological example by performing both model selection and parameter estimation to experimental data gathered from the cardiac Na^+^/K^+^ ATPase.

The Na^+^/K^+^ ATPase is an active ion transporter that pumps three Na^+^ ions out of the cell in exchange for the transport of two K^+^ ions into the cell. As both of these processes are against the typical electrochemical gradients within cells, a source of energy is required to drive this process: ATP hydrolysis. Therefore the reaction catalysed by this enzyme is

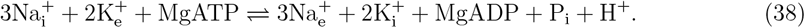

Because ions are transported across a charged membrane, there is interplay between chemical and electrical energy in this process. An advantage of the bond graph representation is that it is general enough to represent both chemical and electrical processes, making it possible to model electrogenic enzymes such as the Na^+^/K^+^ ATPase [44]. Further detail on the incorporation of the membrane potential and thermodynamic constraints are described in Appendix C of the electronic supplementary material.

The Na^+^/K^+^ ATPase is believed to act via the Post-Albers mechanism [52] that is used in many existing models [10, 44, 53]. Here, we generate alternative mechanisms of the Na^+^/K^+^ ATPase based on the Post-Albers mechanism by assigning individual binding and unbinding steps to reactions within enzyme cycles while preserving the overall reaction. Alternative models of the Na^+^/K^+^ ATPase can be generated by assigning individual reactants and products to various binding and unbinding steps in different orders. To examine the effects of incorporating known mechanistic information, we employ two model generation strategies: the “random” (or “unordered”) and “ordered” strategies. In the random strategy, binding steps are assigned in a random order. In the ordered strategy, we only sample from the subset of models in which the binding steps are ordered according to the Post-Albers cycle (see Appendix C of the electronic supplementary material for further detail). Thus, the random sampling strategy may represent a situation where we are performing model development in the absence of prior mechanistic information, whereas the ordered sampling strategy is a representation of a situation where model development is informed by known mechanistic information. We only present results for the ordered strategy here, with fits for the random strategy in the electronic supplementary material (Figures S5–S6). Because the number of possible mechanisms of the Na^+^/K^+^ ATPase are too numerous for an exhaustive search to be computationally feasible, we limit our analysis to a maximum of 500 models for each number of states.

We plot the minimised objective function (defined in Eq. 32) in Figure 5A, with models sampled using the ordered strategy. As expected, due to the larger model space, the objective decreases when considering models of greater complexity. However, the benefit of increasing complexity tapers off after six states, suggesting that relatively simple models may be sufficient to explain the data provided. In models with four or more states, there is a substantial proportion of models that fit particularly poorly, indicating that models may need to satisfy certain structural constraints to explain the experimental data.

**Figure 5:**
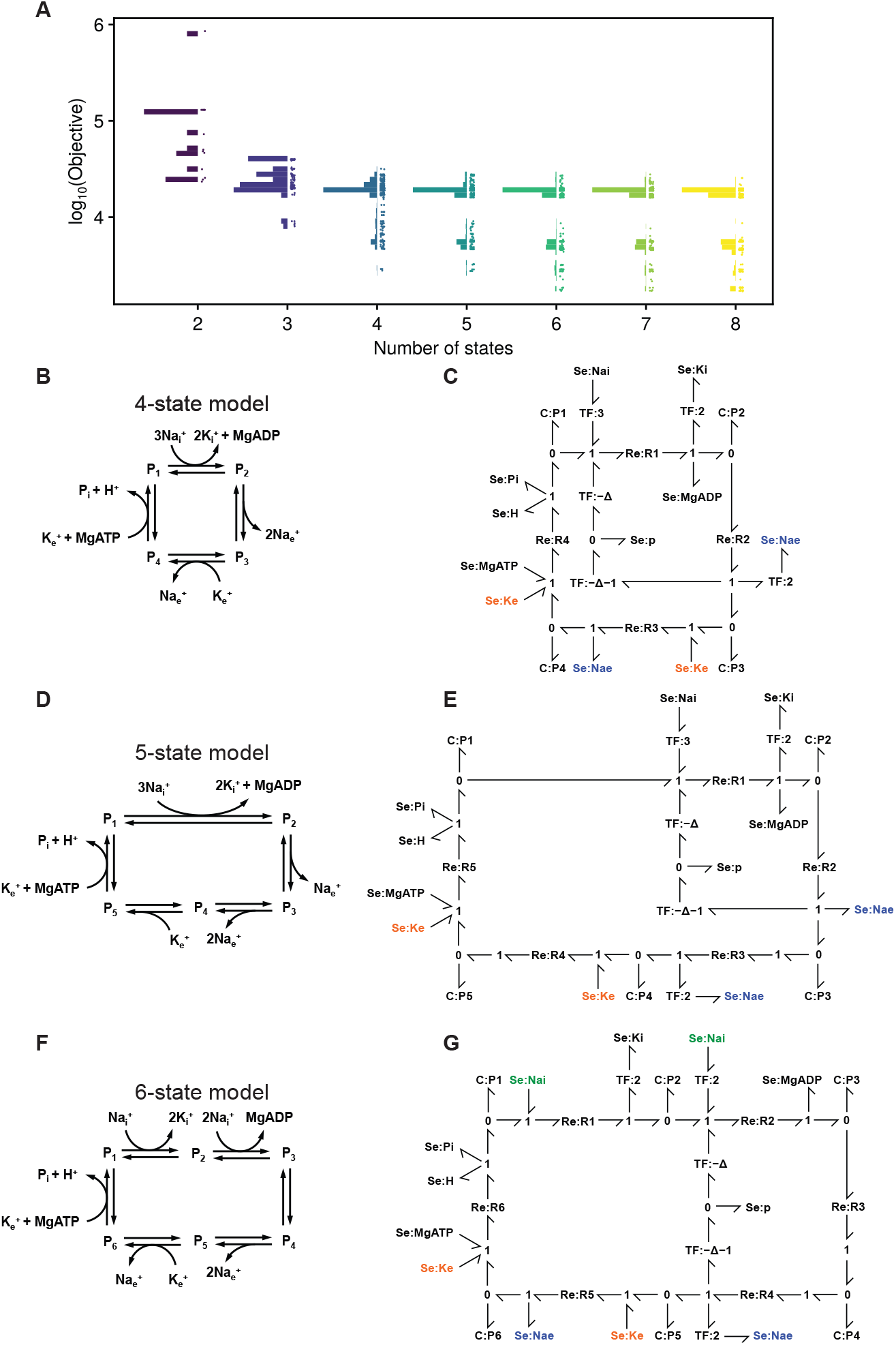
Fits of possible models of the Na^+^/K^+^ ATPase to data. (A) For each number of states, up to 500 models are sampled. Models are sampled with binding steps ordered using the Post-Albers scheme. The objective function is plotted on the vertical axis, with models grouped by the number of reactions. Dots indicate individual models, with the distribution of minimised objectives summarised by histograms to the left of the dots. The best-performing models for 4–6 states are depicted in (B), (D) and (E) respectively, with corresponding bond graphs in (C), (E) and (G) to the right of each reaction network. **Se** components with the same name (emphasised with colour) are connections to the same **Se** component through a common **0** junction [26, 34].

The best-performing models are shown in Figure 5B,D,F, with corresponding bond graphs in Figure 5C,E,G. Interestingly, despite the different number of reactions, the reaction schemes share similarities in network structure. Notably, these networks favour grouping MgADP unbinding with intracellular 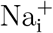 binding, and all networks contained a reaction of the form 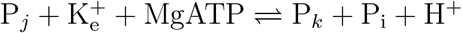. The fits of these models are very comparable (Figure 6). All models provide good fits to voltage-dependence data (Figure 6A–B), extracellular potassium (Figure 6D) and MgATP dependence (Figure 6E). The four- and five-state models are unable to capture the behaviour of the transporter at low intracellular sodium concentrations (Figure 6C); this could be an issue in physiological simulations where intracellular Na^+^ is expected to be low. However, this issue is resolved when moving to the six state model. There is little benefit gained from increasing the complexity of the model beyond six states in terms of fitting to the data. Acknowledging that only a small subset of the full model space was tested, we tentatively suggest that the six-state model provides the best balance between model simplicity and degree of match to data.

**Figure 6:**
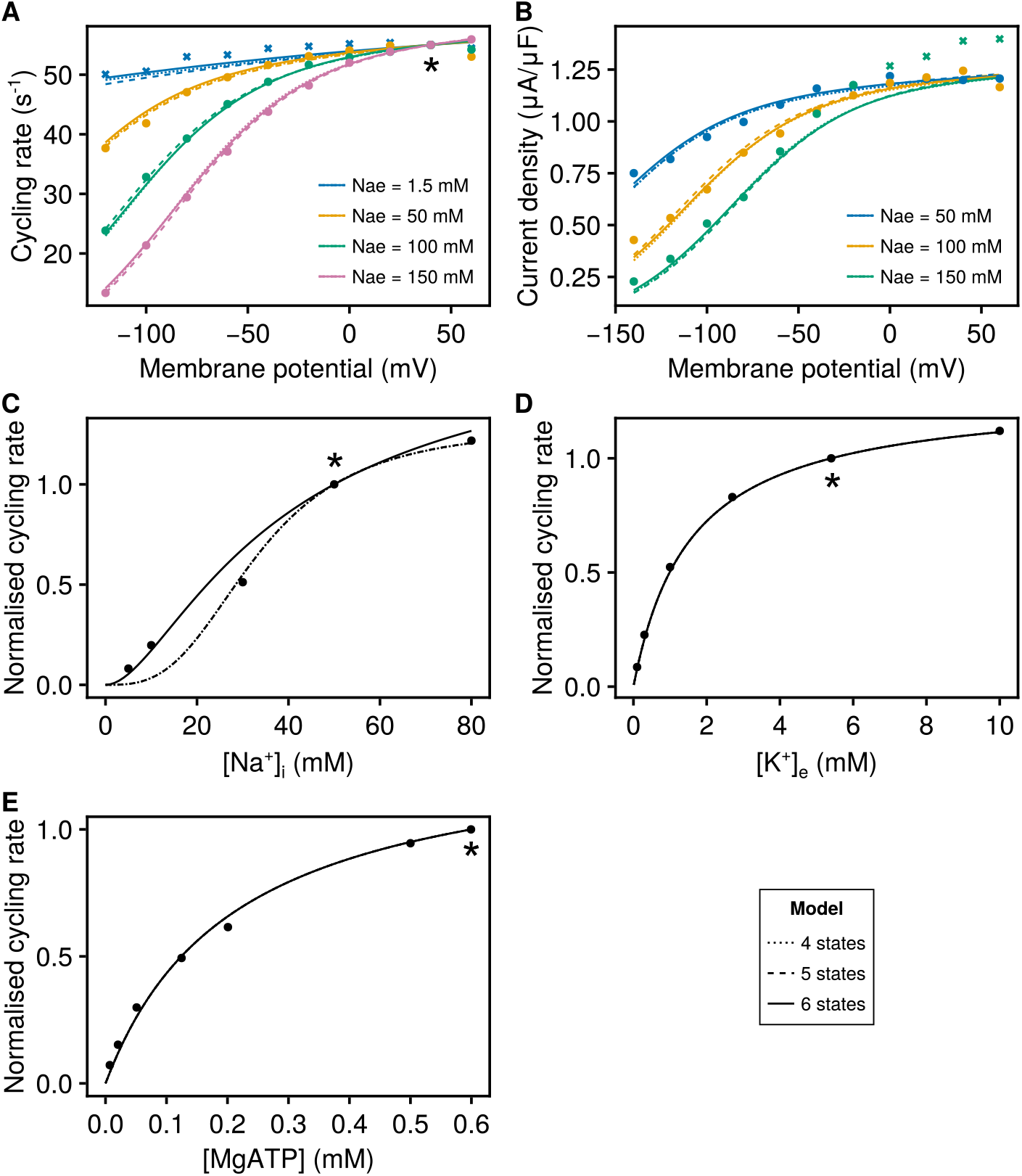
Fits of selected Na^+^/K^+^ ATPase mechanisms to data. Model simulations are compared to **(A-B)** extracellular Na^+^ data (Figs. 3 and 2A of [49]) containing cycling rate (left) and current density (right) measurements, with points and lines grouped by colour corresponding to extracellular Na^+^ concentration, dots representing data used in calibration and crosses data excluded from calibration; **(C)** intracellular Na^+^ data (Fig. 7A of [54]); **(D)** extracellular K^+^ data (Fig. 11A of [49]); **(E)** ATP dependence data (Fig. 3B of [55]). Simulations are performed under the conditions specified in Table 2 and *T* = 310 K. Where present, the asterisks (*) indicate normalisation points. The models refer to the best-performing models in Figure 5B–G. Model outputs from the 4– and 5–state models overlap with each other in (C), and outputs overlap for all plotted models in (D) and (E).

## 4 Discussion

In this study, we introduce an approach to developing models of enzyme cycles for whole-cell modelling. Two key features of our approach are the automatic generation of rate laws and the use of an energetic framework. We develop a methodology for automatically deriving simplified rate laws for a wide range of enzyme cycles with arbitrary complexity, allowing modellers to test multiple models against experimental measurements without the need to manually derive rate laws. In contrast to kinetic parameters, energetic parameters provide a natural basis for incorporating thermodynamic constraints and additional data that may further refine the parameters [16, 42, 44]. We encode information within the energetic formulation using bond graphs, a framework that explicitly models energy and therefore preserves links to fundamental thermodynamic and mechanistic information when models are simplified. The approach is applied to the Na^+^/K^+^ ATPase, generating simple, thermodynamically consistent models that fit the training data well while making minimal assumptions about the mechanism of the enzyme. Our approach could form part of a workflow to automatically build and update components within a whole-cell model.

**Table 2:**
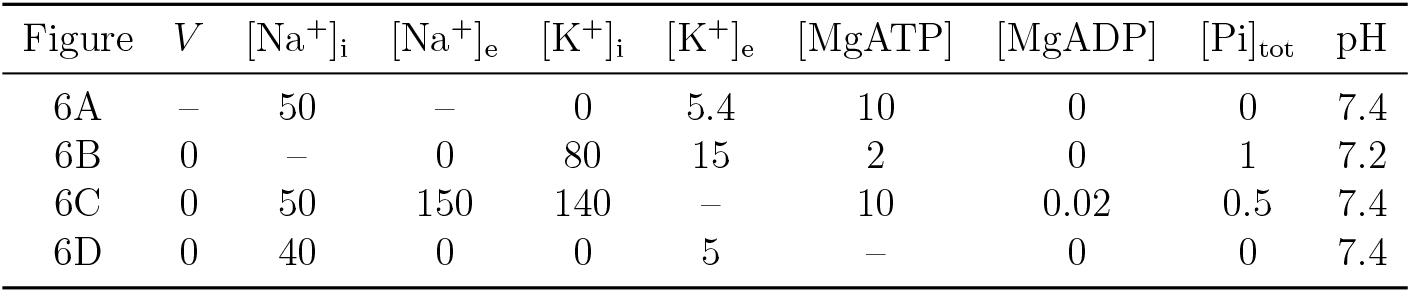
Simulation conditions for Figure 6. Voltages (*V* ) have unit mV, concentrations have unit mM and the pH is dimensionless. Empty entries indicate that the variable was experimentally varied.

| Figure | $V$ | $[\text{Na}^+]_i$ | $[\text{Na}^+]_e$ | $[\text{K}^+]_i$ | $[\text{K}^+]_e$ | $[\text{MgATP}]$ | $[\text{MgADP}]$ | $[\text{Pi}]_{\text{tot}}$ | pH |
| --- | --- | --- | --- | --- | --- | --- | --- | --- | --- |
| 6A | – | 50 | – | 0 | 5.4 | 10 | 0 | 0 | 7.4 |
| 6B | 0 | – | 0 | 80 | 15 | 2 | 0 | 1 | 7.2 |
| 6C | 0 | 50 | 150 | 140 | – | 10 | 0.02 | 0.5 | 7.4 |
| 6D | 0 | 40 | 0 | 0 | 5 | – | 0 | 0 | 7.4 |

While there are numerous methods for simplifying biochemical systems [56, 57], the link between the underlying biochemical mechanism and mathematical equations is often non-existent or lost in the process of simplification [58, 59] and few approaches account for thermodynamic constraints [10, 15]. We therefore base our method on existing approaches that use the King-Altman-Hill method [10, 13, 37, 40]. We improve on these approaches in two ways: by using energetic parameters to automatically satisfy thermodynamic constraints, and by deriving an explicit form for the reduced model so that the method is easily programmable. We note that while the King-Altman-Hill method has been implemented computationally [60], these implementations do not enforce thermodynamic consistency and do not readily extend to multi-physics systems.

Our model calibration suggests that models ranging from three to six states are sufficient to explain steady-state measurements collected from the Na^+^/K^+^ ATPase, depending on whether the sampled models are restricted to the consensus Post-Albers mechanism. The selected models are simpler than many other models that have been used to explain the same data [44, 53]. However, there may be biological phenomena outside the training data that can only be captured by biochemically detailed models. Despite this, these results demonstrate the utility of our approach in model development as it is flexible enough to be adapted depending on the prior mechanistic information that is available, such as the order of binding.

We believe that bond graphs are ideal for whole-cell modelling because their modularity allows full models of enzymes to be interchanged with reduced representations. In particular, bond graphs support the switching of detailed models containing full dynamical behaviour with simplified modules that contain only a rate law [26, 34, 36]. These reduced components are dissipative, thermodynamically consistent, and reusable between different enzyme catalysed reactions. Liebermeister, Uhlendorf and Klipp [61] argue for developing a relatively small library of thermodynamically consistent rate laws that can approximately represent the kinetics of most enzymes. We envisage that simplified rate laws could be used to form such a library, containing relatively simple mechanisms such as the Michaelis-Menten, ping-pong and bi-bi mechanisms. These can be used as first choices to fit new data from enzymes, with further refinements possible when experimental data become comprehensive enough to require more complex mechanisms.

Our approach facilitates the use of model selection, which could form the basis of a method for identifying the correct mechanism of an enzyme in the future. We have described a procedure that ranks models based on their consistency with the data, substantially reducing the space of plausible models. For instance, the models of Na^+^/K^+^ ATPase within the upper band seen in Figure 5A could be easily rejected by this approach. Our results suggest that the steady-state measurements from enzyme assays are generally insufficient to perfectly resolve the mechanism of an enzyme, even in the presence of noiseless and comprehensive data (Figure 4D–F). Therefore, additional information from transient measurements, time series measurements and structural data would likely be required to confidently infer the correct mechanism [50, 62, 63]. Because our simplified models are explicitly linked to a full biophysical mechanism with a system of differential equations, it is relatively easy to extend our approach to model identification to account for time-dependent behaviour. Furthermore, our analysis has the potential to inform subsequent experiments to maximise the mechanistic information that could be gained. In this context, it may be useful to use existing mathematical analyses for mechanism distinguishability [50] as well as Bayesian inference to quantify mechanistic uncertainty [20, 64] while accounting for prior knowledge of the model structure.

A limitation of our approach is the assumption that the concentrations of enzymes are small so that they would be in quasi-steady-state. However, there are studies that have cast doubt over the validity of this assumption [65]. Surovtsova et al. [66] use computational singular perturbation to assess the validity of rapid equilibrium and quasi-steady-state approximations. While their approach does not incorporate thermodynamic information, integrating such a method into a physical and modular framework may allow simulation software to automatically switch between full and reduced models as needed.

## 5 Conclusion

We develop a generalised approach for model reduction based on bond graphs, an energetic framework. Our approach is applicable to a wide range of enzyme cycles, allowing numerous models to be simultaneously tested against experimental measurements. Because our analysis is based on a physical and energy-based framework, the reduced models are thermodynamically consistent and retain links to the underlying biochemical mechanism. As a result, it is relatively straightforward to incorporate thermodynamic information into the development of enzyme models and to reconcile measurements from different experiments. Therefore, our approach contributes to the automation of model development for enzymes while providing an explicit physical framework for coupling models together.

### Software

The model reduction method described in this manuscript is implemented in a Julia package available at https://github.com/mic-pan/EnzymeModelIdentification.jl. The code for generating figures in this manuscript can be found at https://github.com/mic-pan/bond_graph_model_reduction and is archived on Zenodo at https://doi.org/10.5281/zenodo.20551917 [67].

## Dedication

The authors would like to dedicate this study to the late Edmund Crampin, who played a pivotal role in introducing bond graphs to systems biology. Edmund was a kind and valued colleague with an unwavering enthusiasm for his research and helped shape the careers of many, including the authors of this manuscript.

## Acknowledgements

M.P. would like to acknowledge financial support provided by an Australian Government Research Training Program Scholarship. P.J.G. would like to thank the Melbourne School of Engineering for its support via a Professorial Fellowship. This research was in part funded by the Australian Research Council Centre of Excellence for the Mathematical Analysis of Cellular Systems (project number CE230100001).

