## Supplementary Material for "A general thermodynamic approach for model reduction of enzyme cycles and electrogenic transporters"

Michael Pan, Peter J. Gawthrop, Joseph Cursons, Edmund J. Crampin

---

**D Supplementary figures** **10**

### A Model reduction

#### A.1 Summary

The model reduction approach involves finding an explicit mathematical expression for the steady-state cycling rate of an enzyme cycle after it has been reduced using rapid equilibrium. An outline of the procedure is as follows:

Step 1: Separate the system into fast and slow subsystems (Subsection A.3)
Step 2: Analyse the system on the fast timescale to find the rapid equilibrium constraints (Subsection A.4)
Step 3: Find an expression for the rates of change for the slow variables on the slow timescale (Eq. S12)
Step 4: Using the mapping between enzyme states and slow variables from Step 3, derive a linear differential equation for the enzyme states (Eq. S17)
Step 5: Derive the steady-state and conservation constraints from the differential equation on the slow subsystem (Subsection A.5)
Step 6: Derive expressions for the enzyme states at the steady state of the slow subsystem (Eq. S20)
Step 7: Calculate the steady-state reaction rates using the steady-state enzyme states (Eq. S25)
Step 8: Find the cycling rate of the enzyme (Eq. S28)

#### A.2 Notation

The stoichiometry of a network of biochemical reactions is denoted by the forward and reverse matrices $N^f$ and $N^r$, which have rows corresponding to species and columns corresponding to reactions. The stoichiometry of the $i$th species on the forward side of the $j$th reaction is recorded in the $(i, j)$ entry in $N^f$, and a similar convention follows for $N^r$. The stoichiometric matrix is given by $N = N^r - N^f$. In the upcoming analysis, it is useful to break these stoichiometric matrices into partitions. We denote such partitions as $N_\alpha^\beta$ where $\alpha \in \{F, S\}$ and $\beta \in \{\text{cd}, \text{cs}\}$. The symbol $\alpha$ indicates that the matrix is limited to reactions operating on the fast ($F$) or slow ($S$) timescale, and $\beta$ indicates that the matrix is limited to species corresponding to metabolites (chemostatic, or cs) or enzyme states (chemodynamic, or cd). If a symbol is not present, all reactions or species are included. A similar convention applies to $N_\alpha^{f,\beta}$ and $N_\alpha^{r,\beta}$.

We define $\kappa$ and $K$ as vectors containing the reaction and species energetic parameters respectively. The bold symbols $\boldsymbol{\kappa}$ and $\mathbf{K}$ denote diagonal matrices containing these parameters. Using a similar notation to stoichiometric matrices, $\kappa_\alpha$, $K_\beta$, $\boldsymbol{\kappa}_\alpha$, $\mathbf{K}^\beta$ and $X^\beta$ represent vectors and matrices restricted to a subset of species and reactions.

$I_{n\times n}$ is the $n \times n$ identity matrix, and $0_{m\times n}$ is the $m \times n$ zero matrix. The functions $\mathbf{Exp}$ and $\mathbf{Ln}$ are the element-wise exponential and logarithm respectively, and $\circ$ is the element-wise multiplication operator.

#### A.3 Timescale separation

In models of enzyme kinetics, we often assume that some reactions operate on timescales far quicker than the rest. We can depict a vector bond graph of the full biochemical network in Figure S1. The components are represented in italics, denoting the fact that they contain multiple potentials and flows. In the context of a whole-cell model, we see the reactions at rapid equilibrium as occurring on a "fast" timescale, the evolution of enzyme kinetics occurring on a "slow" timescale and the evolution of metabolites occurring on an "ultraslow" timescale [1].

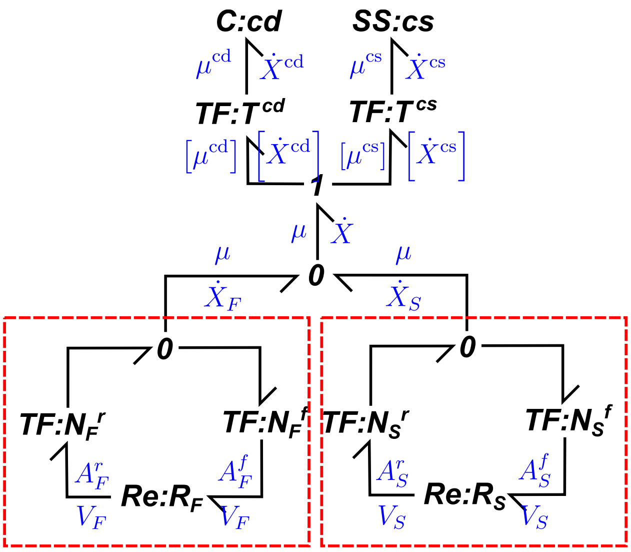

**Figure S1: Generalised model of full enzyme kinetics.**

Since the full model contains both dynamic species and chemostats, they have been placed in separate vector components. The transformers described by matrices $T^{\mathrm{cd}}$ and $T^{\mathrm{cs}}$ are used to merge the chemical potentials into a single vector. To merge the $n_{\mathrm{cd}}$ dynamic species and $n_{\mathrm{cs}}$ chemostats into a vector of length $n_s = n_{\mathrm{cd}} + n_{\mathrm{cs}}$, we define $T^{\mathrm{cd}}$, a $n_s \times n_{\mathrm{cd}}$ logical matrix with 1's where the species match. The $n_s \times n_{\mathrm{cs}}$ matrix $T^{\mathrm{cs}}$ is defined similarly. As a result, the effort coming out of the 1 junction in Figure S1 can be written as

$$\mu = T^{\mathrm{cd}}\mu^{\mathrm{cd}} + T^{\mathrm{cs}}\mu^{\mathrm{cs}}. \tag{S1}$$

As shown in Gawthrop and Crampin [2], the equations of the full model can be described by

$$\dot{X}^{\mathrm{cd}} = N_{\mathrm{cd}}V, \tag{S2}$$

where $X^{\mathrm{cd}}$ is a vector containing the amounts of the enzyme states and $V$ is a vector of the reaction rates. We then split $V$ into $V_S$ and $V_F$, which are the components corresponding to the fast and slow reactions respectively. Then

$$\begin{aligned}\dot{X}^{\mathrm{cd}} &= N_F^{\mathrm{cd}}V_F + N_S^{\mathrm{cd}}V_S && \text{(S3)}\\ &= N_F^{\mathrm{cd}}\boldsymbol{\kappa}_F\left(\mathbf{Exp}\left[{N_F^f}^T\mathbf{Ln}(\mathbf{K}X)\right] - \mathbf{Exp}\left[{N_F^r}^T\mathbf{Ln}(\mathbf{K}X)\right]\right)\\ &\quad + N_S^{\mathrm{cd}}\boldsymbol{\kappa}_S\left(\mathbf{Exp}\left[{N_S^f}^T\mathbf{Ln}(\mathbf{K}X)\right] - \mathbf{Exp}\left[{N_S^r}^T\mathbf{Ln}(\mathbf{K}X)\right]\right). && \text{(S4)}\end{aligned}$$

We also split the chemical potentials into those for the chemodynamic species, and the chemostats, so that

$$\begin{aligned}\dot{X}^{\mathrm{cd}} = N_F^{\mathrm{cd}}\boldsymbol{\kappa}_F&\left(B_F^f\mathbf{Exp}\left[{N_F^{f,\mathrm{cd}}}^T\mathbf{Ln}(\mathbf{K}^{\mathrm{cd}}X^{\mathrm{cd}})\right] - B_F^r\mathbf{Exp}\left[{N_F^{r,\mathrm{cd}}}^T\mathbf{Ln}(\mathbf{K}^{\mathrm{cd}}X^{\mathrm{cd}})\right]\right)\\ &+ N_S^{\mathrm{cd}}\boldsymbol{\kappa}_S\left(B_S^f\mathbf{Exp}\left[{N_S^{f,\mathrm{cd}}}^T\mathbf{Ln}(\mathbf{K}^{\mathrm{cd}}X^{\mathrm{cd}})\right] - B_S^r\mathbf{Exp}\left[{N_S^{r,\mathrm{cd}}}^T\mathbf{Ln}(\mathbf{K}^{\mathrm{cd}}X^{\mathrm{cd}})\right]\right),\end{aligned} \tag{S5}$$

where $\mathbf{K}^{\mathrm{cd}}$ is a diagonal matrix with entries corresponding to the chemodynamic species. For $a \in \{S, F\}$ and $d \in \{f, r\}$, we define the chemostatic contribution as

$$B_a^d = \mathrm{diag}\left(\mathbf{Exp}\left[{N_a^{d,\mathrm{cs}}}^T\mathbf{Ln}(\mathbf{K}^{\mathrm{cs}}X^{\mathrm{cs}})\right]\right). \tag{S6}$$

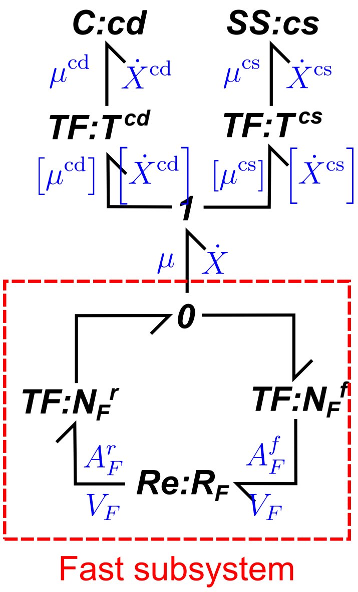

**Figure S2: Fast timescale of enzyme kinetic models.**

In models of enzymes, all reactions are transitions between enzyme states, therefore $N^f_{\text{cd}}$ and $N^r_{\text{cd}}$ contain a single 1 in each column, and zeros elsewhere. As a result, the model is a linear system with the equations

$$\dot{X}^{\text{cd}} = \left[ N_F^{\text{cd}} \boldsymbol{\kappa}_F \left( B_F^f N_F^{f,\text{cd}^T} - B_F^r N_F^{r,\text{cd}^T} \right) + N_S^{\text{cd}} \boldsymbol{\kappa}_S \left( B_S^f N_S^{f,\text{cd}^T} - B_S^r N_S^{r,\text{cd}^T} \right) \right] \mathbf{K}^{\text{cd}} X^{\text{cd}}. \tag{S7}$$

In the next sections, we separate the fast and slow timescales, and show how this separation of timescales can be used to reduce the system of equations.

#### A.4 Fast timescale

The fast timescale contains the reactions that proceed at faster kinetics. We write $\boldsymbol{\kappa}_F = (1/\varepsilon)\boldsymbol{\kappa}_F^*$, and observe the behaviour of the system as $\varepsilon \to 0$. On the fast timescale $T = t/\varepsilon$, Eq. S7 becomes

$$\frac{dX^{\text{cd}}}{dT} = N_F^{\text{cd}} \left[ \boldsymbol{\kappa}_F^* \left( B_F^f N_F^{f,\text{cd}^T} - B_F^r N_F^{r,\text{cd}^T} \right) + \varepsilon \boldsymbol{\kappa}_S \left( B_S^f N_S^{f,\text{cd}^T} - B_S^r N_S^{r,\text{cd}^T} \right) \right] \mathbf{K}^{\text{cd}} X^{\text{cd}}. \tag{S8}$$

Then in the limit as $\varepsilon \to 0$, the rates of the slow reactions vanish, thus

$$\frac{dX^{\text{cd}}}{dT} = N_F^{\text{cd}} V_F^{\text{cd}} = N_F^{\text{cd}} \boldsymbol{\kappa}_F^* \left( B_F^f N_F^{f,\text{cd}^T} - B_F^r N_F^{r,\text{cd}^T} \right) \mathbf{K}^{\text{cd}} X^{\text{cd}}. \tag{S9}$$

This equation corresponds to the reduced bond graph in Figure S2, where the slow subsystem has been removed.

In timescale separation, the steady state of the fast timescale is coupled to the slow dynamics. While steady states in biochemical networks are not necessarily in equilibrium, we make the additional assumption that the steady state is in equilibrium on the fast timescale (otherwise fluxes would be drawn from the chemostats at arbitrarily high rates). This leads to the *rapid equilibrium* constraint

$$\left( B_F^f N_F^{f,\text{cd}^T} - B_F^r N_F^{r,\text{cd}^T} \right) \mathbf{K}^{\text{cd}} X^{\text{cd}} = 0, \tag{S10}$$

which will be used on the slow timescale. As with rapid equilibrium approximations in general, this does not imply that $V_F^{\text{cd}} = 0$, rather that it is determined by the dynamics of the slower subsystem.

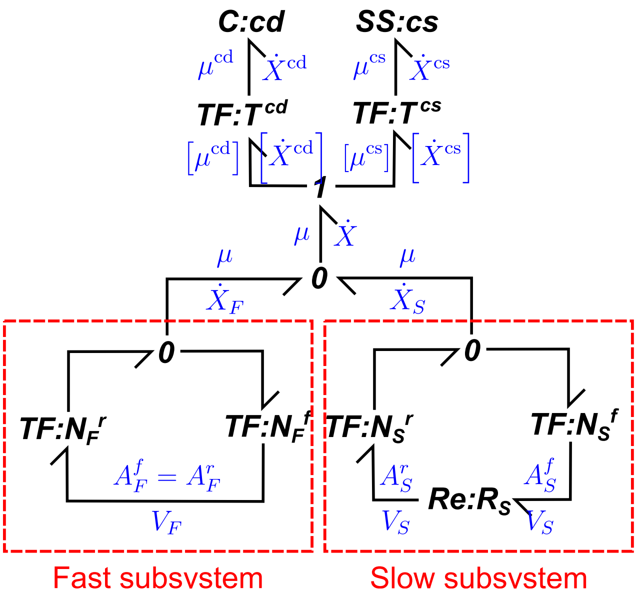

**Figure S3:** **Slow timescale of enzyme kinetic models.**

#### A.5 Slow timescale

If we take $\varepsilon \to 0$, then on the slow timescale, the fast reactions will have run to equilibrium (i.e. Eq. S10 holds). A bond graph representation of this reduced system is shown in Figure S3. To model rapid equilibrium, the **Re** components corresponding to the fast reactions have been removed, so that the forward and reverse affinities are equal [3].

Because many of the states in $X^{\mathrm{cd}}$ are fast variables in the limit $\varepsilon \to 0$, we formulate the differential equations of the slow timescale in terms of the slow variables $Z$. As seen in Schauer and Heinrich [4], the slow variables can be defined as $Z = G_F X^{\mathrm{cd}}$, where $G_F$ is the left nullspace matrix of $N_F^{\mathrm{cd}}$. Then the rates of change of the slow variables are

$$\dot{Z} = G_F \left[ N_F^{\mathrm{cd}} \boldsymbol{\kappa}_F \left( B_F^f N_F^{f,\mathrm{cd}^T} - B_F^r N_F^{r,\mathrm{cd}^T} \right) + N_S^{\mathrm{cd}} \boldsymbol{\kappa}_S \left( B_S^f N_S^{f,\mathrm{cd}^T} - B_S^r N_S^{r,\mathrm{cd}^T} \right) \right] \mathbf{K}^{\mathrm{cd}} X^{\mathrm{cd}} \tag{S11}$$

$$= G_F N_S^{\mathrm{cd}} \boldsymbol{\kappa}_S \left( B_S^f N_S^{f,\mathrm{cd}^T} - B_S^r N_S^{r,\mathrm{cd}^T} \right) \mathbf{K}^{\mathrm{cd}} X^{\mathrm{cd}} \tag{S12}$$

because $G_F N_F^{\mathrm{cd}} = 0$. Because the equality holds regardless of the value of $\varepsilon$, the equation also holds in the limit as $\varepsilon \to 0$.

In order to express Eq. S12 as a proper ODE, we convert between $X^{\mathrm{cd}}$ and $Z$. On the slow timescale, this is possible because the rapid equilibrium constraint (Eq. S10) and $Z = G_F X^{\mathrm{cd}}$ imply

$$\begin{bmatrix} 0_{n_F,1} \\ Z \end{bmatrix} = C X^{\mathrm{cd}}, \tag{S13}$$

where

$$C = \begin{bmatrix} \left( B_F^f N_F^{f,\mathrm{cd}^T} - B_F^r N_F^{r,\mathrm{cd}^T} \right) \mathbf{K}^{\mathrm{cd}} \\ G_F \end{bmatrix}. \tag{S14}$$

If the fast subsystem contains no reaction cycles, $C$ is square and invertible because its rows and columns can be rearranged to form a block diagonal matrix with invertible subblocks. Then

$$X^{\mathrm{cd}} = C^{-1} \begin{bmatrix} 0_{n_F,1} \\ Z \end{bmatrix} = E_Z Z, \tag{S15}$$

where $n_F$ is the number of fast reactions and $E_Z$ is a matrix of the final $n_{\mathrm{cd}} - n_F$ rows of $C^{-1}$.

Using Eq. S15, we can write a differential equation for the slow variables:

$$\dot{Z} = G_F N_S^{\mathrm{cd}} \boldsymbol{\kappa}_S \left( B_S^f N_S^{f,\mathrm{cd}^T} - B_S^r N_S^{r,\mathrm{cd}^T} \right) \mathbf{K}^{\mathrm{cd}} E_Z Z. \tag{S16}$$

We can also write an ODE for all of the chemodynamic species:

$$\dot{X}^{\mathrm{cd}} = E_Z G_F N_S^{\mathrm{cd}} \boldsymbol{\kappa}_S \left( B_S^f N_S^{f,\mathrm{cd}^T} - B_S^r N_S^{r,\mathrm{cd}^T} \right) \mathbf{K}^{\mathrm{cd}} X^{\mathrm{cd}}. \tag{S17}$$

At steady state, the system follows the *quasi-steady-state* constraint

$$\dot{Z} = G_F N_S^{\mathrm{cd}} \boldsymbol{\kappa}_S \left( B_S^f N_S^{f,\mathrm{cd}^T} - B_S^r N_S^{r,\mathrm{cd}^T} \right) \mathbf{K}^{\mathrm{cd}} X^{\mathrm{cd}} = 0. \tag{S18}$$

The final constraint dictating the steady state relates to conserved moieties within the enzyme cycle. These can be summarised using $G$, the left nullspace matrix of $N^{\mathrm{cd}}$ so that $G\dot{X}^{\mathrm{cd}} = GNN^{\mathrm{cd}}X^{\mathrm{cd}} = 0$. Then the steady state must satisfy

$$GX^{\mathrm{cd}} = GX^{\mathrm{cd}}(0). \tag{S19}$$

Eq. S19 can be seen as *conservation relations* determined by the initial conditions of the system. If there is only a single enzyme, $G$ is a row vector of ones.

##### A.6 Reduced equations

We use the following three constraints to derive expressions for $X^{\mathrm{cd}}$ at steady state:

1. The rapid equilibrium constraint in Eq. S10
2. The quasi-steady-state constraint in Eq. S18
3. The conservation constraints in Eq. S19

Since these constraints are all linear functions of $X^{\mathrm{cd}}$, we can combine them into a matrix equation

$$MX^{\mathrm{cd}} = b, \tag{S20}$$

where

$$M = \begin{bmatrix} \left( B_F^f N_F^{f,\mathrm{cd}^T} - B_F^r N_F^{r,\mathrm{cd}^T} \right) \mathbf{K}^{\mathrm{cd}} \\ G_F N_S^{\mathrm{cd}} \boldsymbol{\kappa}_S \left( B_S^f N_S^{f,\mathrm{cd}^T} - B_S^r N_S^{r,\mathrm{cd}^T} \right) \mathbf{K}^{\mathrm{cd}} \\ G \end{bmatrix} \tag{S21}$$

and

$$b = \begin{bmatrix} 0_{n_F \times 1} \\ 0_{n \times 1} \\ GX^{\mathrm{cd}}(0) \end{bmatrix}. \tag{S22}$$

We have defined $n_F$ as the number of fast reactions, and $n$ as the number of slow variables. Eq. S20 can be solved to find an expression for $X^{\mathrm{cd}}$.

Using the expression for $X^{\mathrm{cd}}$, the expressions for the slow reactions $V_S$ can be found using the equation

$$V_S = \boldsymbol{\kappa}_S \left( B_S^f N_S^{f,\mathrm{cd}^T} - B_S^r N_S^{r,\mathrm{cd}^T} \right) \mathbf{K}^{\mathrm{cd}} X^{\mathrm{cd}}. \tag{S23}$$

At steady state, $N^{\mathrm{cd}}V = 0$. By splitting this equation into fast and slow components,

$$N_S^{\mathrm{cd}}V_S + N_F^{\mathrm{cd}}V_F = 0, \tag{S24}$$

therefore the rates of the fast reactions can be found by solving the linear equation

$$N_F^{\mathrm{cd}}V_F = -N_S^{\mathrm{cd}}V_S. \tag{S25}$$

Thus the rates of all reactions at steady state can be found by joining $V_S$ and $V_F$:

$$V = T_S V_S + T_F V_F, \tag{S26}$$

where $T_S$ and $T_F$ are matrices that ensure the fast and slow reactions are mapped to the correct entries. The molar flow rates supplied by the chemostats are given by

$$V_{\mathrm{cs}} = N^{\mathrm{cs}}V. \tag{S27}$$

$V_{\mathrm{cs}}$ can be used to determine the fluxes to and from each metabolite when incorporated into a larger model.

From Gawthrop and Crampin [5], the steady-state cycling rates can be reduced by pathway analysis. By computing the right nullspace matrix $C_p$ of $N^{\mathrm{cd}}$, we find that the steady-state reaction rates can be reduced using the equation

$$V = C_p V_p, \tag{S28}$$

where $V_p$ is a reduced vector with length equal to the number of cycles, and contains the information required to reconstruct $V$. While enzymatic mechanisms in general may have many cycles [5, 6], the examples we deal with in this study only have a single cycle, so $V_p = v$ contains only a single entry that we refer to as the steady-state flux within the main text. $V_p$ can be normalised by $e_0$ to obtain the cycling rate. In the general case where there are multiple pathways, $V_p$ can be solved using the equation

$$V_p = C_p^+ V, \tag{S29}$$

where $C_p^+$ is the Moore-Penrose pseudoinverse of $C_p$.

### B Fitting energetic parameters to kinetic data

Energetic parameters are related to kinetic parameters by the equation

$$\mathbf{Ln}(\mathbf{k}) = \mathbf{M}\mathbf{Ln}(\boldsymbol{\lambda}) \tag{S30}$$

with

$$\mathbf{k} = \begin{bmatrix} k^+ \\ k^- \end{bmatrix}, \quad \mathbf{M} = \left[\begin{array}{c|c} I_{n_r \times n_r} & {N^f}^T \\ \hline I_{n_r \times n_r} & {N^r}^T \end{array}\right], \quad \boldsymbol{\lambda} = \begin{bmatrix} \kappa \\ K \end{bmatrix} \tag{S31}$$

where $k^+$ is the vector of forward kinetic rate constants and $k^-$ is the vector of reverse rate constants [7, 8]. The mapping between kinetic and energetic parameters is not one-to-one. As a consequence of this, more than one set of energetic parameters map to the same set of kinetic parameters [8]. Because steady-state measurements depend only on the kinetics of the system and not the thermodynamic quantities specifically, there will be families of energetic parameters with the same kinetic properties, and therefore explain steady-state data equally well. Accordingly, we designed our parameter estimation method to restrict the space of sampled parameters to those with unique kinetic behaviour and output results that indicate the extent to which energetic parameters are determined.

If there are energetic parameters that are known prior to parameter estimation, Eq. S30 can be expressed as

$$\mathbf{Ln}(\mathbf{k}) = \mathbf{M}_k\mathbf{Ln}(\boldsymbol{\lambda}_k) + \mathbf{M}_u\mathbf{Ln}(\boldsymbol{\lambda}_u), \tag{S32}$$

where $\mathbf{M}_k$ and $\mathbf{M}_u$ are the columns of $\mathbf{M}$ corresponding to the known and unknown parameters respectively. Similarly, $\boldsymbol{\lambda}_k$ and $\boldsymbol{\lambda}_u$ are the rows of $\boldsymbol{\lambda}$ corresponding to the known and unknown parameters respectively. The unknown parameters can then be found by solving the equation

$$l = \mathbf{M}_u\mathbf{Ln}(\boldsymbol{\lambda}_u), \tag{S33}$$

where $l = \mathbf{Ln}(\mathbf{k}) - \mathbf{M}_k\mathbf{Ln}(\boldsymbol{\lambda}_k)$. In many cases, even after previously known energetic parameters are set, the mapping between kinetic and remaining energetic parameters is still non-unique. To overcome this issue, we choose energetic parameters to set prior to parameter estimation. Energetic parameters associated with linearly dependent rows of $\mathbf{M}_u$ are arbitrarily set until a one-to-one mapping results. The resulting set of energetic parameters can then be generated from this individual point.

For any two vectors $\boldsymbol{\lambda}'_u$ and $\boldsymbol{\lambda}''_u$ that satisfy Eq. S33,

$$l = \mathbf{M}_u\mathbf{Ln}(\boldsymbol{\lambda}'_u) = \mathbf{M}_u\mathbf{Ln}(\boldsymbol{\lambda}''_u) \tag{S34}$$

and therefore

$$\mathbf{M}_u[\mathbf{Ln}(\boldsymbol{\lambda}''_u) - \mathbf{Ln}(\boldsymbol{\lambda}'_u)] = 0. \tag{S35}$$

Once a set of unknown parameters $\boldsymbol{\lambda}'_u$ arises from fitting, all other possible parameters follow the form

$$\boldsymbol{\lambda}''_u = \mathbf{Exp}(\mathbf{r}) \circ \boldsymbol{\lambda}'_u, \tag{S36}$$

where $\mathbf{r} \in \ker(\mathbf{M}_u)$. If $R$ is a right nullspace matrix of $\mathbf{M}_u$, then the family of possible energetic parameters is described by the set

$$\mathcal{S} = \left\{\boldsymbol{\lambda}''_u = \mathbf{Exp}(\mathbf{t}R) \circ \boldsymbol{\lambda}'_u : \mathbf{t} \in \mathbb{R}^{1\times n_{\text{free}}}\right\}. \tag{S37}$$

In the above equation, $\mathbf{t} = [t_1 \quad t_2 \quad \ldots \quad t_{n_{\text{free}}}]$ is a row vector containing the free parameters, with a length equal to the dimension of $\ker(\mathbf{M}_u)$.

### C Fitting experimental data for Na⁺/K⁺ ATPase

#### C.1 Incorporation of membrane potential

Unlike many enzymes, the $Na^+/K^+$ ATPase transports ions across a charged membrane. Therefore, the cycling rate must be dependent on membrane potential in order to be thermodynamically consistent. Bond graphs are highly desirable in this context because they can represent quantities from different domains under a single framework. Under the bond graph representation, chemical potential ($\mu$) and membrane potential ($V_m$) are analogous quantities. We make use of this

**Table S1: The binding steps used to generate candidate models for the $Na^+/K^+$ ATPase.** Due to the cyclic nature of enzyme cycles, if $n$ is equal to the number of states, $P_1$ appears in the products rather than $P_{n+1}$. The species p represents the translocation of a single unit of charge from the intracellular to the extracellular side of the membrane, and $\Delta$ is a partitioning constant.

| Step # | Reaction |
|---|---|
| 1 | $P_n \rightleftharpoons P_{n+1} + K_i^+$ |
| 2 | $P_n \rightleftharpoons P_{n+1} + K_i^+$ |
| 3 | $P_n + Na_i^+ \rightleftharpoons P_{n+1}$ |
| 4 | $P_n + Na_i^+ \rightleftharpoons P_{n+1}$ |
| 5 | $P_n + Na_i^+ - \Delta p \rightleftharpoons P_{n+1}$ |
| 6 | $P_n \rightleftharpoons P_{n+1} + MgADP$ |
| 7 | $P_n \rightleftharpoons P_{n+1} + Na_e^+ - (1+\Delta)p$ |
| 8 | $P_n \rightleftharpoons P_{n+1} + Na_e^+$ |
| 9 | $P_n \rightleftharpoons P_{n+1} + Na_e^+$ |
| 10 | $P_n + K_e^+ \rightleftharpoons P_{n+1}$ |
| 11 | $P_n + K_e^+ \rightleftharpoons P_{n+1}$ |
| 12 | $P_n \rightleftharpoons P_{n+1} + P_i + H^+$ |
| 13 | $P_n + MgATP \rightleftharpoons P_{n+1}$ |

analogy to incorporate voltage dependence. While $\mathbf{K}^{cs}X^{cs}$ in Eq. S6 contains entries of the form $K_s x_s$, these can also be expressed in terms of the chemical potential:

$$\gamma = K_s x_s = e^{\mu_s/RT}. \tag{S38}$$

We call $\gamma$ the thermokinetic potential [9]. To incorporate voltage dependence, the chemical potential $\mu$ is substituted for the analogous potential $FV_m$, where $F = 96485$ C/mol:

$$\gamma = e^{FV_m/RT}. \tag{S39}$$

#### C.2 Model generation

$Na^+/K^+$ ATPase mechanisms are generated using the 13 binding and unbinding steps listed in Table S1. An unordered mechanism with $n$ states and $n$ reactions is generated by randomly assigning each step to a single reaction. Ordered steps are generated by starting with a cycle with binding steps in the order specified in Table S1, and grouping neighbouring steps together into the same reaction to form the required number of reactions. To avoid intracellular and extracellular species from binding and unbinding in the same reaction, we exclude models that group steps 13 and 1.

#### C.3 Parameter estimation

It is possible to directly incorporate known relationships between energetic parameters into models of enzymes. Since $Na^+$ and $K^+$ have a known equilibrium of equal concentration on the intracellular and extracellular sides of the membrane, we set the energetic constants of these species to $K_{Nai} = K_{Nae} = K_{Ki} = K_{Ke} = 1 mM^{-1}$. To account for the known standard free energy of MgATP hydrolysis of $\Delta^0_{MgATP} = 11.9$ kJ/mol at 310K [10], we set $K_{MgADP} = K_{MgADP} = K_{MgADP} = 1$ mM$^{-1}$ and $K_{MgATP} = 9881$ mM$^{-1}$. Since these species are chemostats, their parameters are expressed in terms of concentration so that they generalise to different compartmental volumes.

Because the concentrations of Pi in Table 2 refer to total Pi rather than free Pi, total Pi is converted to free Pi using the equation

$$[\mathrm{Pi_{free}}] = \frac{[\mathrm{Pi_{tot}}]}{1 + [\mathrm{K_i^+}]/\mathrm{K_{d,KPi}} + [\mathrm{H^+}]/\mathrm{K_{d,HPi}} + [\mathrm{Na_i^+}]/\mathrm{K_{d,NaPi}}}, \quad \text{(S40)}$$

where the dissociation constants are $\mathrm{K_{d,KPi}} = 292$ mM, $\mathrm{K_{d,HPi}} = 10^{-3.77}$ mM and $\mathrm{K_{d,NaPi}} = 224$ mM [11]. While temperature is known to affect enzyme kinetics, this influence has only been characterised phenomenologically from experimental measurements [12]. Due to the difficulty in predicting the precise effect on energetic parameters, we assume a constant temperature of 310 K across all experimental measurements.

#### D Supplementary figures

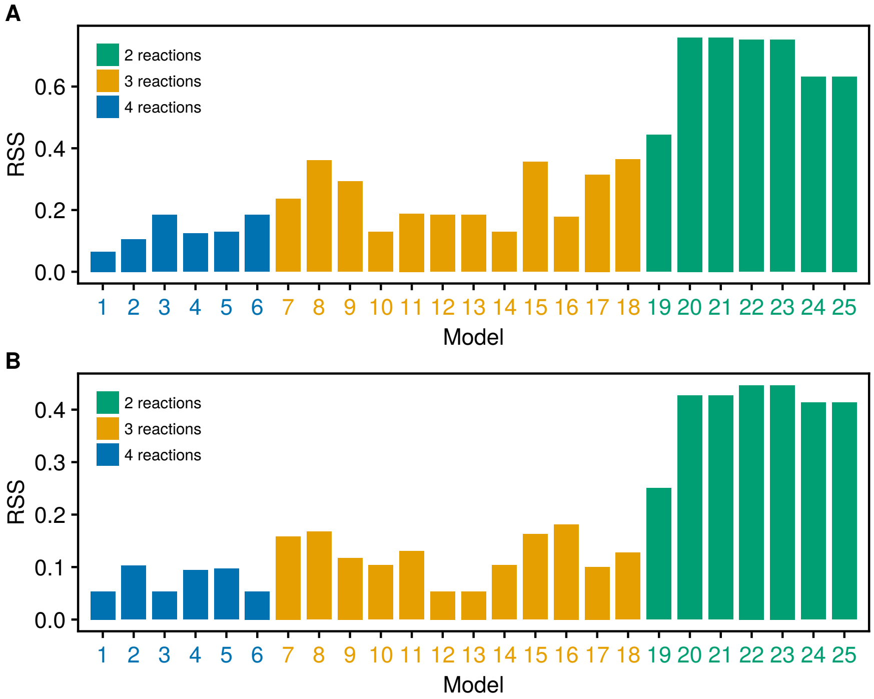

**Figure S4: A comparison of fits of bi-bi enzyme catalytic mechanisms generated from synthetic data with noise.** The residual sum of squares (RSS) resulting from fitting each model to the synthetic data generated from (A) Model 1, and (B) Model 3, with additive Gassian measurement noise with a standard deviation of 0.01.

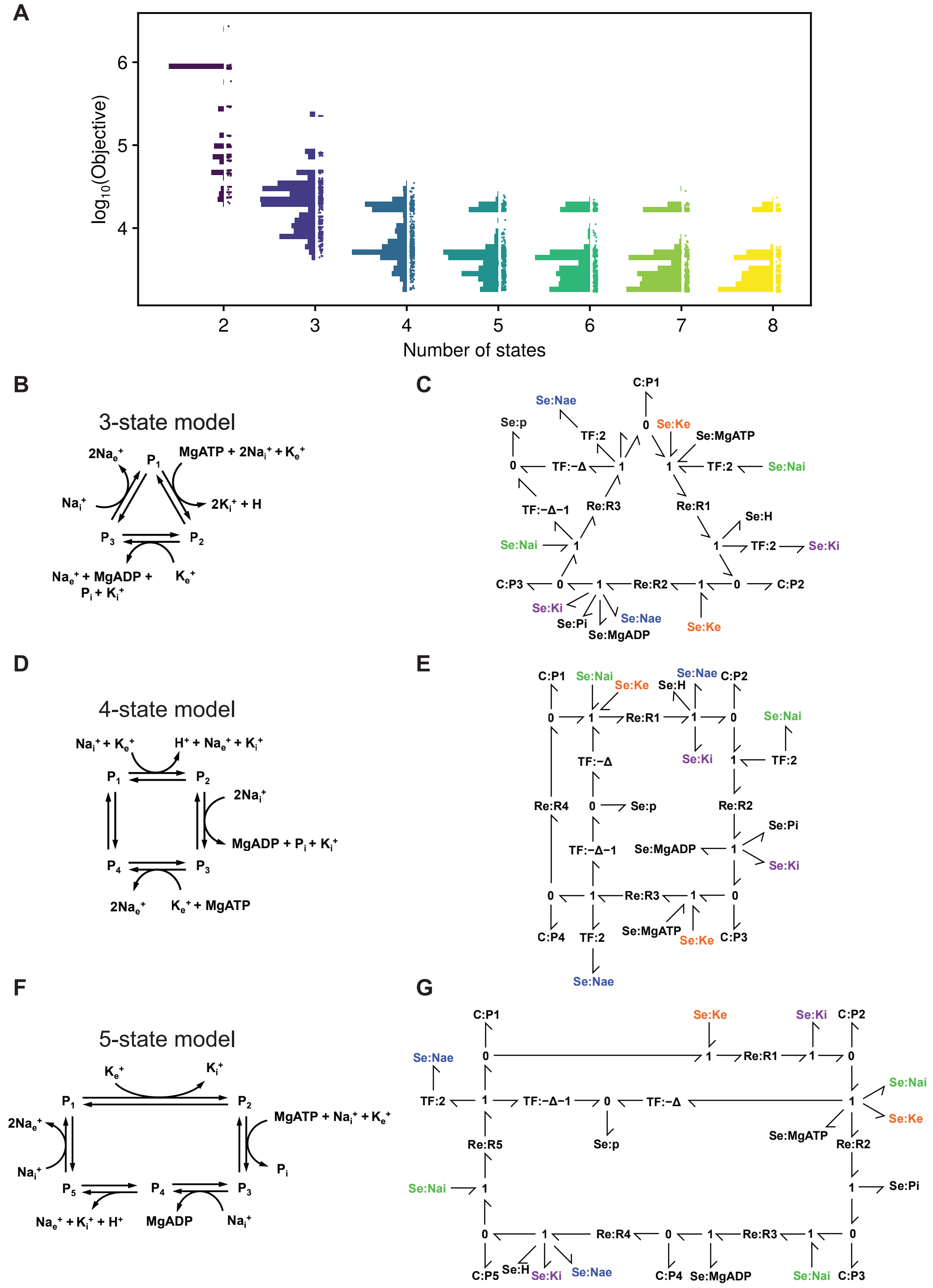

**Figure S5: Fits of random models of the Na$^+$/K$^+$ ATPase to data.** (A) For each number of states, up to 500 models are randomly sampled. The objective function is plotted on the vertical axis, with models grouped by the number of reactions. Dots indicate individual models, with the distribution of minimised objectives summarised by histograms to the left of the dots. The best-performing models for 3–5 states are depicted in (B), (D) and (E) respectively, with corresponding bond graphs in (C), (E) and (G) to the right of each reaction network. **Se** components with the same name (emphasised with colour) are connections to the same **Se** component through a common **0** junction [13, 14].

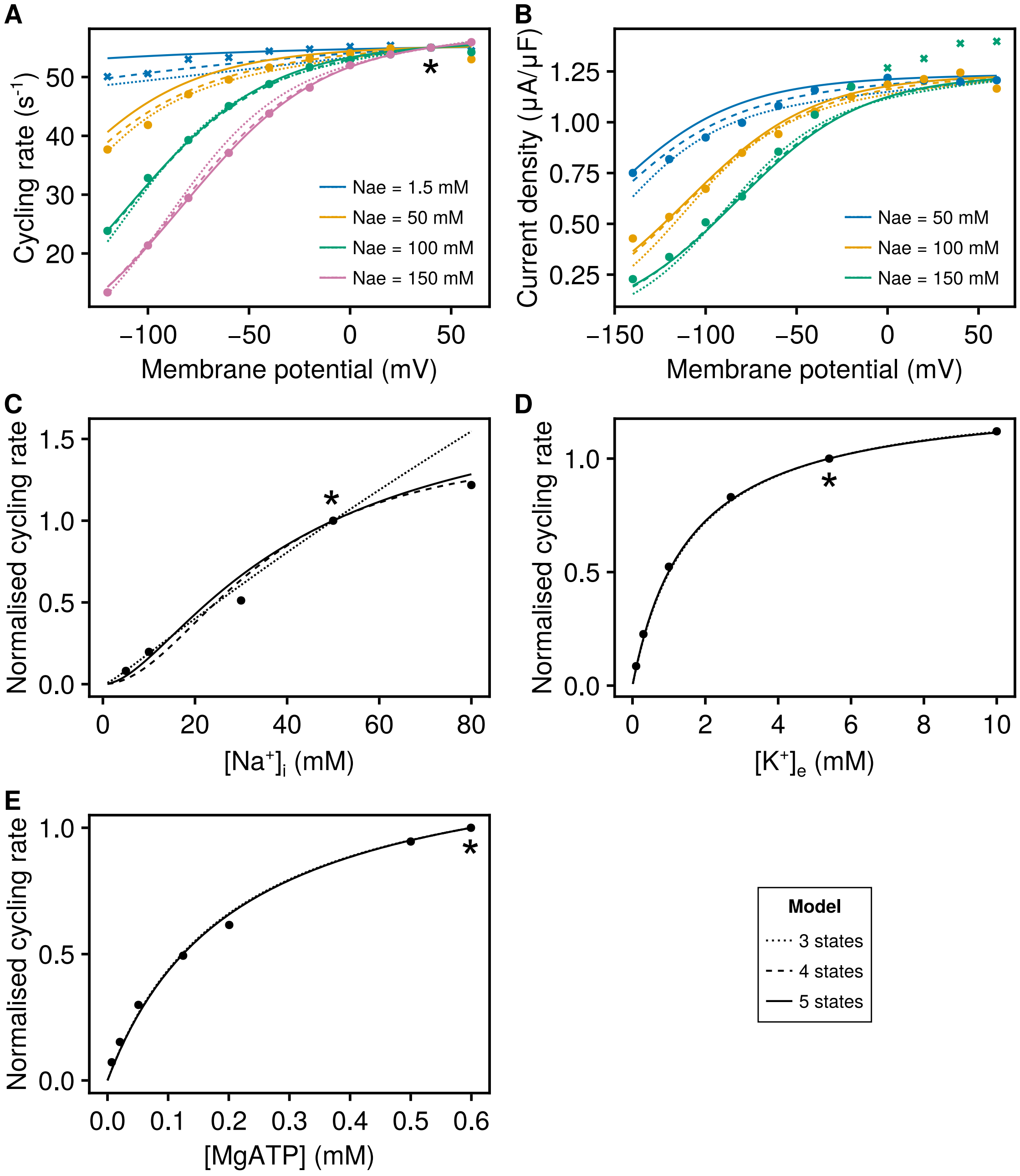

**Figure S6: Fits of selected random $Na^+/K^+$ ATPase mechanisms to data.** Model simulations are compared to **(A-B)** extracellular $Na^+$ data (Figs. 3 and 2A of [15]) containing cycling rate (left) and current density (right) measurements, with points and lines grouped by colour corresponding to extracellular $Na^+$ concentration, dots representing data used in calibration and crosses data excluded from calibration; **(C)** intracellular $Na^+$ data (Fig. 7A of [16]); **(D)** extracellular $K^+$ data (Fig. 11A of [15]); **(E)** ATP dependence data (Fig. 3B of [17]). Simulations are performed under the conditions specified in Table 2 and $T = 310$ K. Where present, the asterisks (*) indicate normalisation points. The models refer to the best-performing models in Figure S5B–G.
